# The mechanotransduction channel Piezo2 refines axonal projections to the accessory optic system and regulates the optokinetic reflex

**DOI:** 10.64898/2026.06.30.735567

**Authors:** Natalie R. Hamilton, Victoria N. Neckles, Timour Al-Khindi, Nitya Donthi, Shun Mizutori, Renata Fu, James K. Kiraly, Bea C. Winship, Alex L. Kolodkin, Karina Chaudhari

## Abstract

The optokinetic reflex (OKR) is an evolutionarily conserved reflexive behavior that ensures image stabilization on the retina during global motion. It consists of smooth eye tracking movements in the direction of the moving stimulus interspersed with rapid resetting saccades. This reflex is driven by retinal ON direction-selective ganglion cells (oDSGCs), which comprise distinct subtypes tuned either to vertical or horizontal motion. oDSGCs convey directional signals to the brain via precise axonal projections to specific accessory optic system (AOS) nuclei. However, the mechanisms that establish and maintain the specificity of these circuits remain poorly understood. Here, we identify a critical role for the mechanosensitive ion channel Piezo2 in refining AOS circuitry to ensure appropriate eye movement responses. Single-cell transcriptomic profiling revealed selective enrichment of Piezo2 in horizontally-tuned oDSGCs. We show that both loss and hyperactivation of Piezo2 in retinal neurons leads to cross-coupling of horizontal and vertical OKR responses, producing aberrant diagonal eye tracking movements during horizontal optokinetic stimulation. Mechanistically, Piezo2 regulates the developmental refinement of oDSGC axonal projections within the AOS, and disruption of this process results in persistent ectopic innervation that enables aberrant crosstalk between horizontal and vertical motion pathways. These findings reveal a channel activity-dependent mechanism that ensures the functional segregation of directional motion circuits underlying gaze stabilization.

**HIGHLIGHTS:**

- *Piezo2* is selectively expressed in forward-tuned ON and ON-OFF DSGCs that innervate the accessory optic system
- Both loss and overactivation of Piezo2 induce cross-coupling of horizontal and vertical optokinetic reflexes
- Piezo2 function in ON-OFF DSGCs is dispensable for normal optokinetic reflex responses
- Retinal Piezo2 refines axonal targeting to the accessory optic system nuclei, ensuring proper optokinetic reflex function

## INTRODUCTION

The accessory optic system (AOS) includes a group of brainstem nuclei that does not participate in image formation but instead drives the optokinetic reflex (OKR) to stabilize gaze during global visual motion^[1–3]^. The OKR consists of slow eye tracking movements (ETMs) in the direction of motion, punctuated by rapid resetting saccades^[1, 4]^. It is an evolutionarily conserved behavior observed across the vertebrate phylogeny, underscoring its fundamental importance for visual stability^[5–7]^. Eye movement disorders including nystagmus affect a significant percentage of the population, and retinal origins for aberrant eye movements have recently been proposed^[8–10]^, highlighting the significance of understanding the mechanisms governing AOS circuit assembly.

The sensory drive for the AOS originates from retinal direction-selective retinal ganglion cells (DSGCs). DSGCs are tuned to respond to motion along a single preferred direction. AOS-projecting DSGCs comprise three subtypes of ON DSGCs (oDSGCs) tuned to slow upward, downward, and forward motion (U-, D-, and F-oDSGCs)^[3, 5, 11]^, and one non-canonical ON-OFF DSGC subtype that responds preferentially to slow forward motion (slow F-ooDSGCs)^[3]^. The functional necessity for two distinct forward-preferring DSGC subtypes, and their relative contributions to the horizontal OKR, remain unclear.

The AOS nuclei controlling vertical and horizontal OKR are anatomically segregated and receive input from correspondingly tuned DSGC subtypes. U- and D-oDSGCs project to the dorsal and ventral subdivisions of the medial terminal nucleus (dMTN and vMTN), respectively, while F-DSGCs (F-oDSGCs and slow F-ooDSGCs) project to the nucleus of the optic tract (NOT) and the dorsal terminal nucleus (DTN)^[2, 3, 12, 13]^. The precise correspondence between DSGC subtype and AOS target nucleus, and the resulting behavioral specificity of the individual AOS nuclei, make this circuit an ideal system for identifying mechanisms governing circuit assembly and refinement. Several molecules controlling the guidance and targeting of AOS DSGCs to their respective AOS nuclei have been identified^[14, 15]^. However, across the CNS axons frequently exhibit broader targeting early in development and circuit pruning is required to achieve precise and selective connectivity^[16–18]^. Whether the AOS exhibits such circuit refinement, and what molecules regulate it, remain largely unknown.

Here, we characterize early misprojections within the AOS that require axonal pruning and refinement to achieve the behavioral segregation of horizontal and vertical OKR. We identify a specific role for the mechanosensitive ion channel Piezo2 in this refinement process, which when abrogated results in a phenotypic cross-coupling of the horizontal and vertical OKR. Further, we establish a functional distinction between F-oDSGCs and slow F-ooDSGCs, demonstrating that Piezo2 function in the latter is dispensable for circuit refinement.

Together, these fundings suggest a novel role for Piezo2 as a critical regulator of F-oDSGC axonal refinement and AOS circuit assembly.

## RESULTS

### Piezo2 is expressed in subsets of retinal ganglion cells and amacrine cells in the mouse retina

To identify molecular regulators of retinal direction-selective (DS) circuits that drive the horizontal OKR, we performed high-depth single-cell RNA sequencing in the *Hoxd10-GFP* transgenic mouse line, which selectively labels AOS-projecting DSGCs^[3]^. In this line, labeled cells include three ON-DSGC (oDSGC) subtypes tuned to upward, downward, or forward motion, as well as an atypical ON-OFF DSGC (ooDSGC) subtype tuned to slow forward motion. GFP^+^ RGCs were sequenced across five developmental stages from E18.5 to P21. Clustering of the filtered data (see Methods) identified three transcriptionally distinct populations (Figure 1A). Notably, these clusters differentially expressed members of the same protein tyrosine phosphatase receptor family: *Ptprk*, *Ptprm* and *Ptprt* (Figure 1B). Our previous work established that *Ptprk* and *Ptprm* are selectively enriched in U-oDSGCs and D-oDSGCs, respectively^[19, 20]^ (Figures 1C and 1D). By exclusion, we inferred that the *Ptprt-*enriched cluster corresponds to forward DSGCs (F-oDSGCs; Figure 1E). Interestingly, the mechanosensitive ion channel *Piezo2* was also strongly enriched in this putative F-DSGC cluster (Figure 1F). Consistent with this result, analysis of a published P5 RGC single-cell dataset^[21]^ also revealed selective co-expression of *Ptprt* and *Piezo2* in the RGC cluster we characterized as F-DSGCs, (Figure 1G).

**Figure 1.**
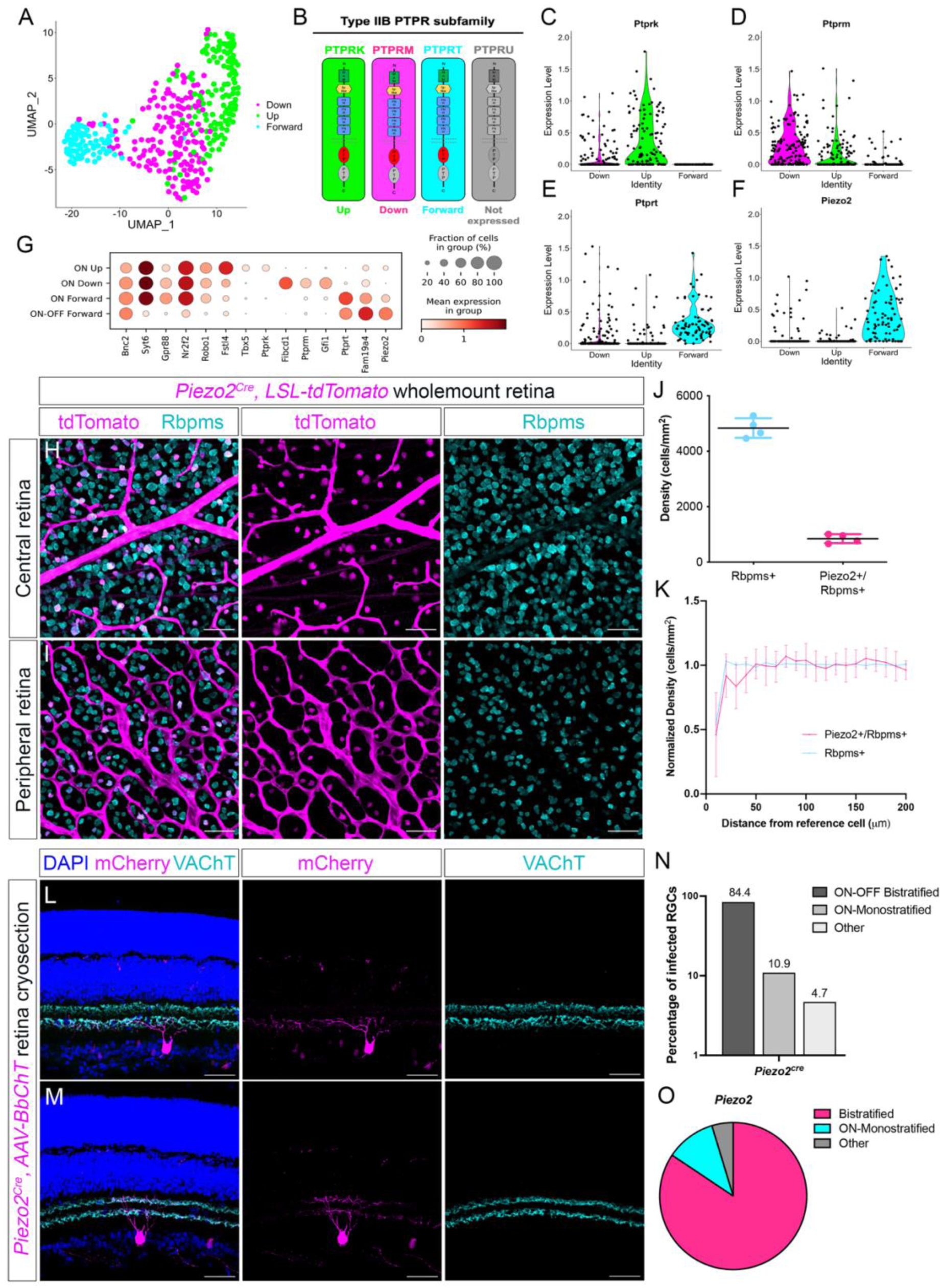
*Piezo2* is expressed in monostratified and bistratified RGCs that co-stratify with SACs. (**A**) Clustering of the filtered *Hoxd10-GFP* dataset revealed three transcriptionally distinct clusters. (**B-E**) These three clusters show complementary expression patterns of three Type IIB PTPRs: one cluster selectively expresses the U-oDSGC marker *Ptprk* **(C)**, another expresses relatively high levels of the D-oDSGC marker *Ptprm* (**D**), and the third, the putative F-DSGC cluster, selectively expresses *Ptprt* (**E**). (**F**) Violin plot showing that *Piezo2* is highly enriched in the putative F-DSGC cluster. (**G**) Dotplot from the P5 RGC scRNA-seq dataset^[21]^ showing enrichment of *Ptprt* and *Piezo2* in *Fam19a4*-expressing perivascular F-ooDSGCs (cluster 29) and in F-oDSGCs (re-clustering of cluster 32). This re-clustering, described in our previous work^[61]^, corroborates *Ptprk* expression in U-oDSGCs and *Ptprm* enrichment in D-oDSGCs. **(H, I**) Wholemount retinas from *Piezo2^Cre^; LSL-tdTomato* adult mice show that *Piezo2* (tdTomato^+^) is expressed in a subset of RGCs (Rbpms^+^, pan RGC marker) throughout the central (**H**) and peripheral (**I**) retina. (**J**) Quantification of RGC and Piezo2^+^ RGC density shows that *Piezo2* is expressed by ∼17% of all RGCs. (**K**) Density Recovery Profile (DRP) analysis reveals a lack of mosaic spacing among Piezo2^+^ RGCs, indicating that this is a heterogeneous population comprising more than one subtype. (**L, M**) RGCs labeled by a Cre-dependent AAV in *Piezo2^Cre^* retinas co-stratify with SAC dendrites (labeled by VAChT) and exhibit either ON-monostratified (**L**) or ON-OFF bistratified (**M**) dendritic morphology. (**N, O**) Quantification of sparse labeling experiments with *Piezo2^Cre^* shows that ∼84% of labeled RGCs are bistratified and ∼11% are monostratified; the “Other” category includes OFF-monostratified and displaced RGCs. Data are presented as mean ± SE. See also Figure S1.

To examine *Piezo2* expression in the mouse retina, we used a *Piezo2^Cre^* mouse line which preserves endogenous Piezo2 function^[22]^. Wholemount immunohistochemistry of *Piezo2^Cre^; LSL-tdTomato* retinas revealed tdTomato expression in Rbpms^+^ retinal ganglion cells (RGCs; Figures 1H and 1I). *Piezo2* is expressed in ∼17% of RGCs, and density recovery profile (DRP) analysis indicated that these cells do not form a mosaic, suggesting that Piezo2 marks multiple RGC subtypes (Figures 1J and 1K). In addition to RGCs, *Piezo2* expression was noted in non-SAC amacrine cell populations (Figures S1A and S1B). To characterize the morphology of Piezo2^+^ RGCs, we performed sparse-labeling of Piezo2^+^ RGCs via intravitreal injection of Cre-dependent AAV-Brainbow-mCherry (*AAV-BbChT*) in P2 *Piezo2^Cre^* mice. We analyzed the morphology of labeled RGCs at P16. Co-labeling with vesicular acetylcholine transporter (VAChT), which marks starburst amacrine cell (SAC) processes, revealed two predominant Piezo2^+^ RGC morphologies that co-stratify with SAC dendrites. Most Piezo2^+^ RGCs displayed ON-OFF bistratified dendrites in S2 and S4 sublayers of the IPL (Figure 1L). This morphology is consistent with that of perivascular slow F-ooDSGCs, which were recently described to express *Piezo2*^[23]^. A smaller population (∼10%) of Piezo2^+^ RGCs exhibited dendrites stratifying primarily in S4, in concordance with Forward oDSGC (F-oDSGC) morphology (Figure 1M). Rare Piezo2^+^ RGCs were observed stratifying in OFF sublayers (Figures 1N and 1O). Similar results were obtained with a genetic sparse-labeling strategy using *Piezo2^Cre^; MORF3* mice. The *MORF3* allele allows for Cre-dependent, sparse, and stochastic expression of a membrane-targeted V5 tag. Piezo2^+^ RGCs expressing the V5 tag also exhibit ON-OFF-bistratified and ON-monostratified morphology, both co-stratifying with SACs (Figures S1C–S1E). Together, these results indicate that *Piezo2* is predominantly expressed in ON-OFF-bistratified and ON-monostratified RGCs, morphologies consistent with AOS F-DSGCs.

### Piezo2 is expressed in F-oDSGCs and F-ooDSGCs that project to the accessory optic system

To determine conclusively whether Piezo2^+^ RGCs correspond to AOS DSGCs, we examined retinas from *Piezo2^Cre^; LSL-tdTomato; Hoxd10-GFP* mice (Figures 2A–2D). Approximately 36% of GFP^+^ DSGCs co-expressed tdTomato (Figure 2E), indicating that *Piezo2* is expressed in a subset of AOS DSGCs, as expected for F-DSGCs. Sparse-labeling of these neurons using *AAV-BbChT* in *Piezo2^Cre^; Hoxd10-GFP* mice confirmed that most double-labeled cells exhibited ON-OFF bistratified morphology, consistent with slow F-ooDSGCs, while a smaller fraction displayed ON-monostratified morphology, consistent with F-oDSGCs (Figure 2F). Reconstructions of sparsely labeled neurons revealed that Piezo2^+^ and Piezo2^-^ *Hoxd10-GFP* DSGCs exhibit similar dendritic arbor morphologies (Figures 2G and 2H). Since morphology alone cannot distinguish between U-, D-, and F-oDSGCs, we relied on markers specific to these three subtypes to corroborate scRNA sequencing data (Figures 1C–1E). We performed *in situ* hybridization on cross-sections of P10 *Piezo2^Cre^; LSL-tdTomato* retinas. While ∼40% of Piezo2^+^ cells in the GCL expressed *Ptprt* (Figure S2A and S2D), they did not express the D-oDSGC marker *Fibcd1* nor the U-oDSGC marker *Ptprk* (Figures S2B–S2D). We analyzed further *Piezo2^Cre^; LSL-tdTomato; Spig1-GFP* retinas, in which the *Spig1-GFP* allele specifically labels U-oDSGCs in the ventronasal retina. No cells co-expressing GFP and tdTomato were observed in the ventronasal retina, confirming that U-oDSGCs do not express *Piezo2*. Thus, using a molecular classification, we conclude that within the AOS, *Piezo2* expression is restricted to horizontally-tuned F-DSGCs.

**Figure 2.**
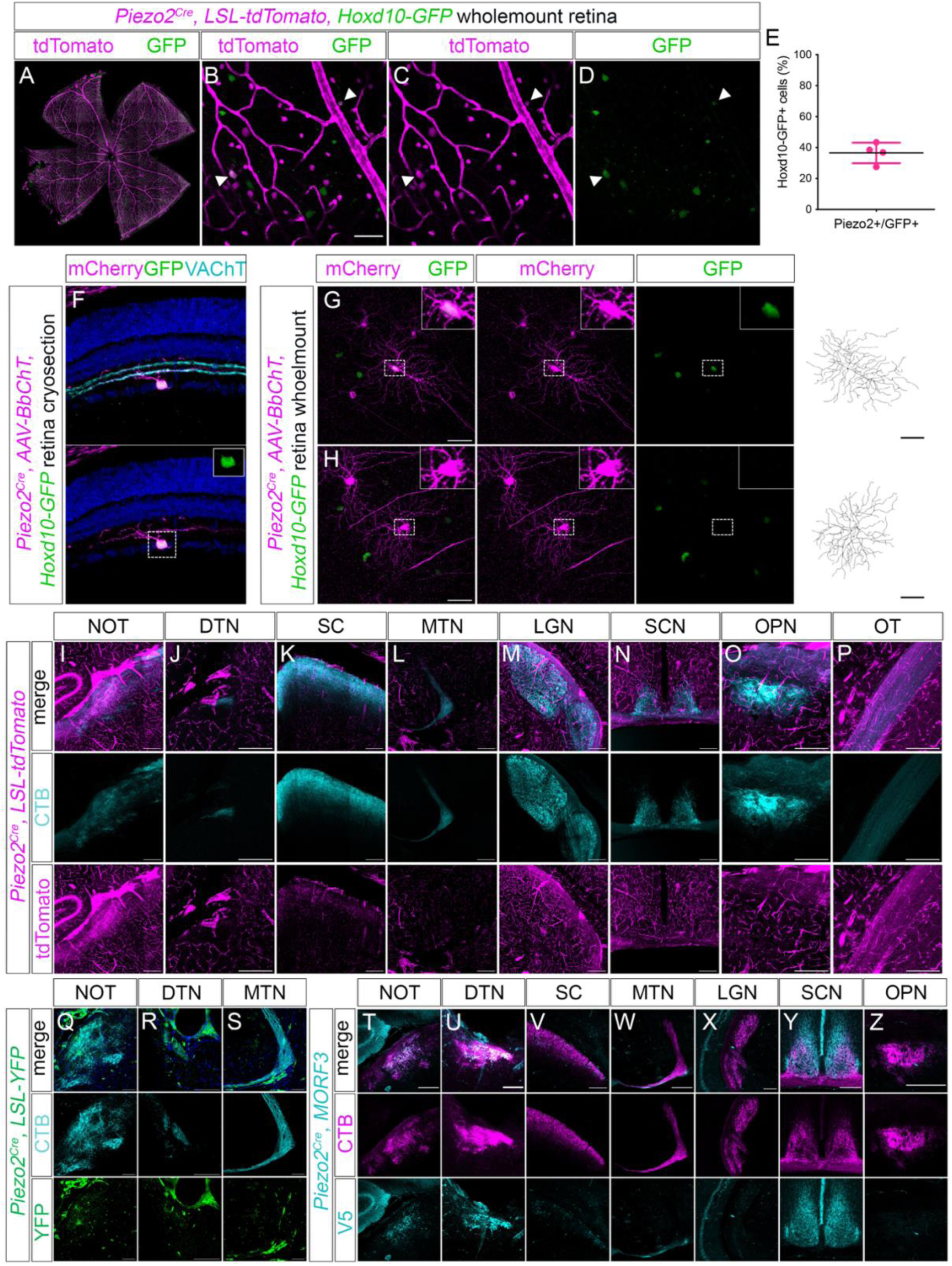
*Piezo2* is expressed in Forward ON and ON-OFF DSGCs that project to the AOS. (**A-D**) Wholemount retinas from *Piezo2^Cre^; LSL-tdTomato; Hoxd10-GFP* mice show that a subset of genetically labeled Piezo2^+^ cells are AOS DSGCs (*Hoxd10-GFP^+^*, white arrowheads). (**E**) Quantification shows that ∼36% of *Hoxd10-GFP* cells express *Piezo2*. (**F**) Sparse-labeling with a Cre-dependent AAV injected intraocular at P2 reveals monostratified Piezo2^+^GFP^+^ RGCs stratifying in S4. (**G, H**) Sparsely labeled Piezo2^+^GFP^+^ and Piezo2^+^GFP^-^ cells have similar dendritic arbors (traces, right). **(I-P**) Retinorecipient nuclei were labeled with bilateral intraocular injections of CTB-647 in adult *Piezo2^Cre^; LSL-tdTomato* mice. *Piezo2* is widely expressed throughout the brain vasculature. Axon terminals are apparent in the NOT (**I**), DTN (**J**) and SC (**K**), but not in the MTN (**L**), LGN (**M**), SCN (**N**), or OPN (**O**). Piezo2^+^ axon terminals can be seen traversing the optic tract (**P**). **(Q-S**) Labeling of retinorecipient nuclei with bilateral intraocular injections of CTB-647 in adult *Piezo2^Cre^; LSL-GFP* mice shows similar axon innervation in the NOT (**Q**) and DTN (**R**), but not the MTN (**S**). **(T-Z**) Labeling of retinorecipient nuclei with bilateral intraocular injections of CTB-555 in adult *Piezo2^Cre^; MORF3* mice, in which stochastic expression of a membrane-targeted V5 tag results in sparse labeling of Piezo2^+^ cells, shows that Piezo2^+^ RGCs innervate the NOT (**T**), DTN (**U**), and SC (**V**), but not the MTN (**W**). Piezo2^+^ RGCs do not appear to innervate the LGN (**X**), SCN (**Y**), or OPN (**Z**), though non-overlapping V5 and CTB-555 staining in the SCN suggests that some SCN neurons may express *Piezo2* independent of retinal innervation (**Z**). Data are presented as mean ± SE. See also Figure S2.

Next, we determined the central targets of Piezo2^+^ RGCs by examining their axonal projections. In the AOS, vertically-tuned oDSGCs project to the medial terminal nucleus (MTN), whereas horizontally-tuned oDSGCs project to the nucleus of the optic tract (NOT) and dorsal terminal nucleus (DTN). Slow F-ooDSGCs have previously been shown to project to the NOT and SC^[3]^. To determine whether Piezo2^+^ RGCs project to these nuclei, we performed intravitreal injection of a fluorescent dye cholera toxin subunit B-Alexa 647 (CTB-647) in adult *Piezo2^Cre^; LSL-tdTomato* and *Piezo2^Cre^; LSL-YFP* mice. Piezo2^+^/tdTomato^+^ axons were observed in the optic tract, NOT and DTN, SC but not the MTN (Figures 2I–2P). Similarly, Piezo2^+^/YFP^+^ axons were observed in the NOT and DTN, but not the MTN (Figures 2Q–2S). However, strong *Piezo2* expression in brain vasculature in these mice limited interpretation of terminal fields in some regions.

To overcome this limitation, we performed CTB-555 injections in adult *Piezo2^Cre^; MORF3* reporter mice, enabling selective labeling of Piezo2^+^ RGC axons via sparser V5 expression. Piezo2^+^ axon terminals were indeed detected in the NOT, DTN and SC but not in the MTN, LGN, SCN, or OPN (Figures 2T–2Z). Strong V5 signal observed within SCN neurons likely reflects endogenous reporter expression rather than retinal input due to non-overlapping V5 and CTB-555 staining patterns (Figure 2Y). Finally, as an alternative method to selectively label Piezo2^+^ RGC axons, we performed intravitreal injections of a Cre-dependent AAV encoding GFP (*AAV2-FLEX-GFP*) in P2 *Piezo2^Cre^* mice. Piezo2^+^ RGC axons appear again to innervate the NOT, DTN, and SC, but not the MTN (Figures S2F–S2M). Taken together, these results strongly suggest that *Piezo2* is expressed in forward-tuned AOS DSGCs projecting to the NOT and DTN, namely F-oDSGCs and slow F-ooDSGCs.

### Piezo2 is required in the retina to prevent cross-coupling of horizontal and vertical optokinetic reflexes responses

Given the selective enrichment of Piezo2 in AOS F-DSGCs, we asked whether Piezo2 contributes to horizontal motion processing. We assessed AOS circuit function by measuring the OKR in head-fixed mice. To selectively remove Piezo2 from the retina, we generated retina-specific *Piezo2* knockout mice (*Ret^Piezo2 LOF^*) using the *Chx10^Cre^* driver, which drives the expression of Cre in retinal progenitor cells^[24]^. *Ret^Piezo2 LOF^* mice displayed normal OKR responses to continuous checkerboard rotation in all four directions – nasal, temporal, upward, and downward (Figures 3A–3D). Eye tracking movements (ETMs) are composed of horizontal (epX) and vertical (epY) components (Figure 3E). During horizontal ETMs, vertical displacement is typically minimal, indicating that the eye moves primarily in the horizontal plane. While this pattern was preserved in both wild type and mutant mice during nasal stimulation, *Ret^Piezo2 LOF^* mice exhibited a prominent vertical component during temporal stimulation, resulting in small but significant diagonal eye movements in phase with temporal ETMs (Figures 3F–3I). Quantification revealed significantly increased upward ETMs during temporal stimulation in *Ret^Piezo2 LOF^* mice, whereas nasal stimuli did not produce this effect (Figures 3J–3M). Importantly, nasal and temporal horizontal gain during sinusoidal stimulation (Figures 3N and 3O) and response latencies to looming stimuli (Figures 3P and 3Q), were unchanged, indicating intact visual function. Thus, retinal Piezo2 is not required for horizontal OKR generation, but is necessary to prevent cross-coupling between horizontal and vertical OKR pathways.

**Figure 3.**
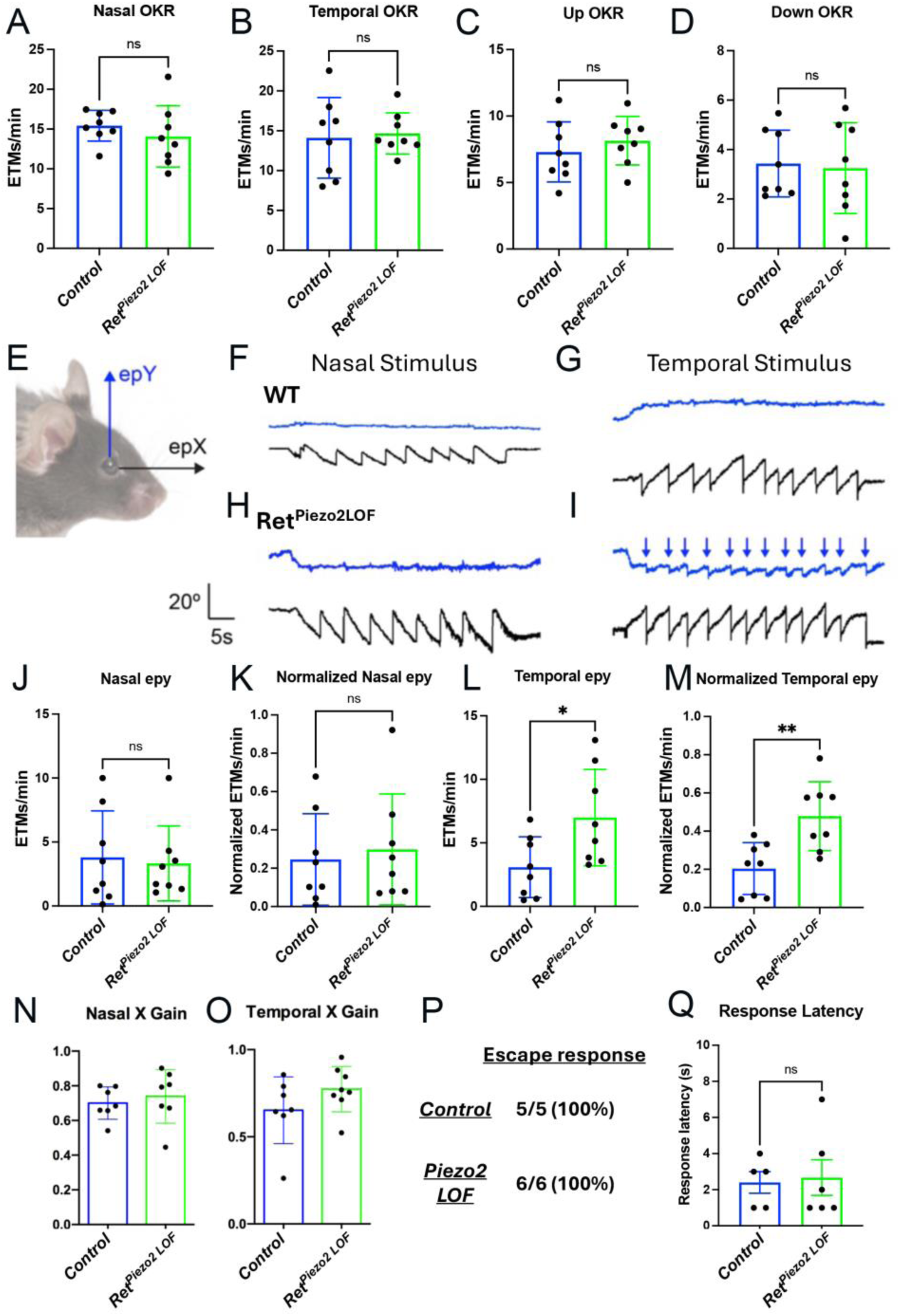
Retina-specific loss of Piezo2 results in cross-coupling of the horizontal and vertical OKR. (**A-D**) Quantification of eye tracking movements (ETMs) per minute in response to continuous gratings moving in the nasal (**A**), temporal (**B**), upward (**C**), or downward (**D**) direction. No statistically significant differences were observed between WT and *Ret^Piezo2LOF^* mice. (**E**) Eye movements are separated into horizontal (epX) and vertical (epY) components. (**F-I**) For nasal stimuli, both WT (**F**) and *Ret^Piezo2LOF^* mice (**H**) exhibit limited vertical eye movements (blue traces) but robust horizontal eye movements (black traces). However, for temporal stimuli, *Ret^Piezo2LOF^* mice (**I**) generate upward eye movements that are in phase with and match the timing of horizontal eye movements (blue arrows), indicating that the eye moves diagonally in response to a temporally moving stimulus. **(J, K**) Absolute and normalized number of in-phase upward vertical ETMs/min (nasal epY) during nasal stimuli. **(L, M**) Absolute and normalized number of in-phase upward vertical ETMs/min (temporal epY) during temporal stimuli; *Ret^Piezo2LOF^* mice make significantly more in-phase upward ETMs in response to temporally-moving stimuli than WT mice. (**N, O**) Gain of eye tracking movements during nasal and temporal stimuli is unchanged in *Ret^Piezo2LOF^* mice. (**P**) *Ret^Piezo2LOF^* mice showed normal escape behavior in the looming task, a measure of the ability to detect overhead looming stimuli. (**Q**) Response latency of WT and *Ret^Piezo2LOF^* mice in the looming assay. Data are presented as mean ± SE. Significance was assessed using Student’s *t*-test. *p < 0.05, **p < 0.01.

### Piezo2 in perivascular slow F-ooDSGCs is dispensable for AOS function

Recent work identified Fam19a4/Nts^+^ perivascular RGCs as a horizontally-tuned ooDSGC subtype that expresses *Piezo2* and regulates retinal vascular patterning, distinct from traditional Cartpt^+^ ooDSGCs^[23]^. To test whether these perivascular RGCs correspond to slow F-ooDSGCs projecting to the AOS, we examined *Nts^Cre^; LSL-tdTomato* retinas. Nts^+^ RGCs comprised ∼7% of total RGCs and were frequently associated with the retinal vasculature, as described previously^.[18]^ (Figures 4A–4C). Notably, crossing this line with *Hox10-GFP* mice revealed that ∼30% of *Hoxd10-GFP* DSGCs co-express tdTomato (Figures 4D and 4E), consistent with the expected fraction of slow F-ooDSGC^[3]^. Meanwhile, double-labeled GFP^+^tdTomato^+^ RGCs comprise a very small fraction of the total Nts^+^ RGCs (Figure 4F), suggesting that either Nts^+^ RGCs comprise two or more molecularly distinct subtypes or that the *Hox10-GFP* transgenic line incompletely labels AOS DSGCs. To differentiate between these possibilities, and to confirm that double-labeled GFP^+^Nts^+^ RGCs are indeed slow F-ooDSGCs and not any of the oDSGC subtypes, we conducted sparse labeling of Nts^+^ RGCs. P2 *AAV-BbChT* injection in *Nts^Cre^* mice revealed that ∼95% of Nts^+^ RGCs exhibit bistratified dendrites in S2 and S4 (Figures 4G–4I), confirming their ON-OFF DSGC identity. These results, coupled with previous work identifying Nts^+^ RGCs as a temporally-preferring (forward in visual space) DSGC subtype,^[23]^ suggest that Nts^+^ RGCs are indeed a homogenous population of ON-OFF bistratified RGCs and that *Hoxd10-GFP* labels only a fraction of all AOS DSGCs. To determine their central targets, we examined axonal projections in *Nts^Cre^* mice following intraocular injection of *AAV-BbChT* and CTB-647. Nts^+^ RGCs projected to the NOT, DTN, and SC, with minimal labeling in the LGN (Figures 4J–4Q). Thus, perivascular Nts^+^ RGCs correspond to slow F-ooDSGCs and target AOS nuclei specialized for horizontal motion.

**Figure 4.**
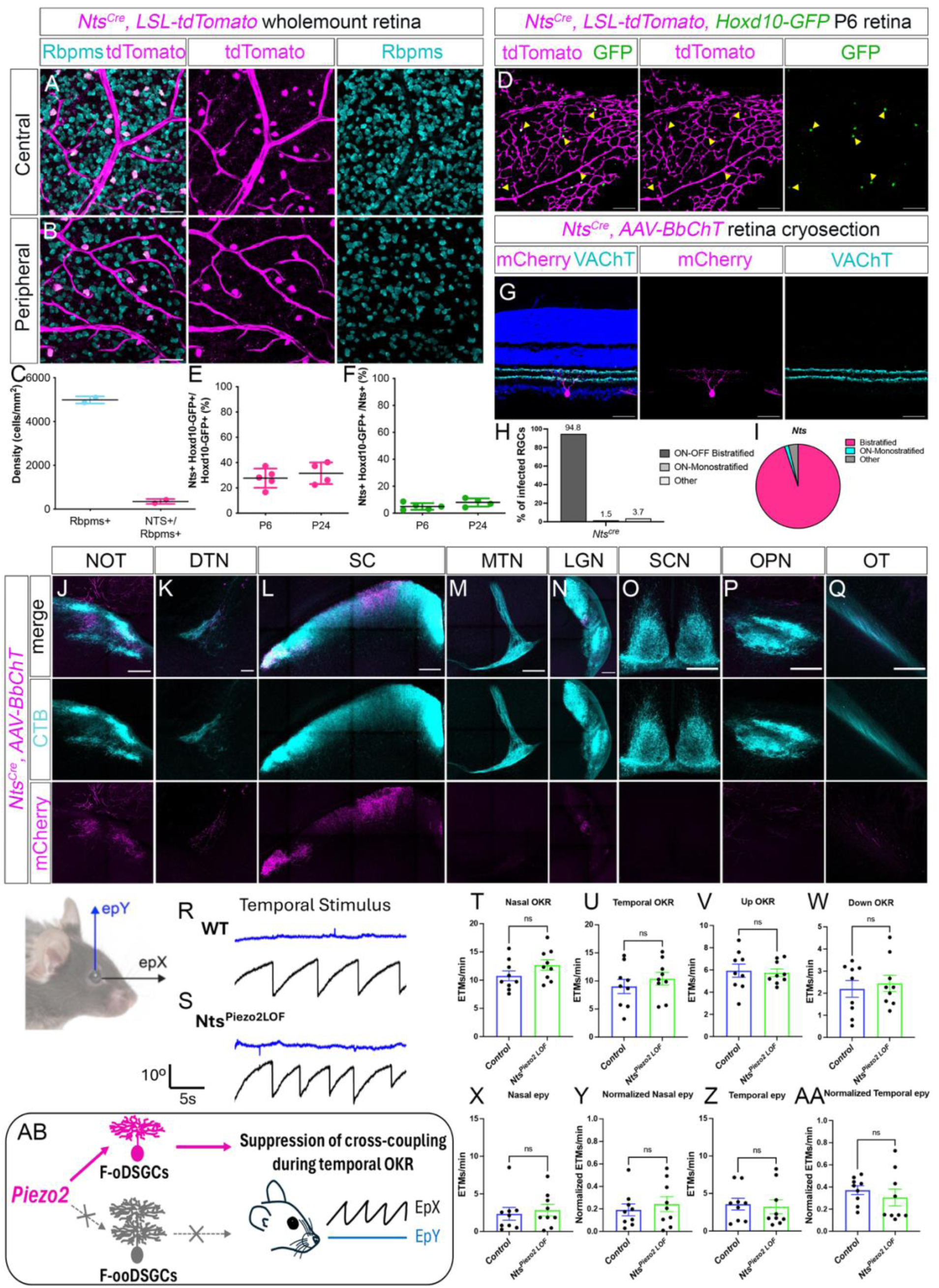
Loss of Piezo2 from perivascular ON-OFF DSGCs does not affect the OKR. **(A, B)** Wholemount retinas from *Nts^Cre^; LSL-tdTomato* adult mice show that *Nts* (tdTomato^+^) is expressed in a subset of RGCs (Rbpms^+^, pan RGC marker) that are frequently located near the vasculature throughout the central (**A**) and peripheral (**B**) retina. (**C**) Quantification of RGC and Nts^+^ RGC density shows that *Nts* is expressed by ∼7% of all RGCs. (**D**) Wholemount retinas from *Nts^Cre^; LSL-tdTomato; Hoxd10-GFP* mice show that a subset of genetically labeled Nts^+^ cells are AOS DSGCs (*Hoxd10-GFP^+^*, yellow arrowheads). (**E, F**) Quantification at P6 and P24 shows that ∼30% of *Hoxd10-GFP* cells express *Nts* (**E**), while only 5-7% of Nts^+^ RGCs express *Hoxd10-GFP* (**F**). (**G**) RGCs labeled by a Cre-dependent AAV in *Nts^Cre^* retinas co-stratify with SAC dendrites (labeled by VAChT) and exhibit primarily ON-OFF bistratified dendritic morphology. (**H, I**) Quantification of sparse labeling experiments with *Nts^Cre^* shows that >94% of labeled RGCs are bistratified; the “Other” category includes OFF-monostratified and displaced RGCs. (**J-Q**) Labeling of axonal projections in *Nts^Cre^* mice following P2 intraocular injections of CTB-647 and a Cre-dependent AAV encoding mCherry shows that Nts^+^ RGCs innervate the NOT (**J**), DTN (**K**), and SC (**L**), but not the MTN (**M**), SCN (**O**), or OPN (**P**); minimal innervation can be seen in the LGN (**N**), and axon terminals can be detected in the optic tract (**Q**). (**R-AA**) Quantification of OKR responses in *Nts^Piezo2LOF^* mice reveals no changes in ETMs per minute in response to continuous gratings moving in the nasal (**T**), temporal (**U**), upward (**V**), or downward (**W**) direction, nor any statistically significant differences in absolute and normalized number of in-phase upward vertical ETMs/min (epY) during nasal (**X, Y**) or temporal (**R, S, Z, AA**) stimuli. (**AB**) Schematic depicting a requirement for Piezo2 in F-oDSGCs, and not F-ooDSGCs, in suppressing cross-coupling of the horizontal and vertical OKR during temporal stimuli. Data are presented as mean ± SE. Significance was assessed using Student’s *t*-test. See also Figure S3.

Because both F-oDSGCs and slow F-ooDSGCs project to the NOT and express *Piezo2*, we tested whether Piezo2 function in slow F-ooDSGCs contributes to OKR circuit function. We generated *Nts^Cre^; Piezo2^Flox^* (*Nts^Piezo2 LOF^*) mice, the same line used previously to achieve effective *Piezo2* deletion from perivascular F-ooDSGCs to demonstrate its requirement for retinal vasculature patterning^[23]^. OKR recordings in *Nts^Piezo2 LOF^* mice revealed normal responses in all four stimulus directions (Figures 4R–4W), and, unlike *Ret^Piezo2 LOF^* mice, these mice showed no increase in upward ETMs during nasal or temporal stimulation (Figures 4X–4AA). Together, these results indicate that Piezo2 function in perivascular slow F-ooDSGCs is dispensable for AOS circuit function (Figure 4AB).

In addition to RGCs, we detected *Piezo2* expression in non-SAC amacrine cell populations (Figures S1A and S1B). To exclude the possibility that Piezo2 function in amacrine cells contributes to the cross-coupling phenotype seen in *Ret^Piezo2 LOF^* mice, we selectively deleted *Piezo2* in amacrine cells using the *Ptf1a^Cre^* driver which specifically expresses Cre in amacrine cell precursors^[25]^. *Ptf1a^Piezo2 LOF^* mice showed normal OKR responses in all four stimulus directions and no evidence of increased cross-coupling in response to horizontally-moving stimuli (Figures S3A–S3J). Collectively, these results indicate that neither amacrine cells nor slow F-ooDSGCs serve as the source for Piezo2 function in suppressing OKR cross-coupling, and they suggest that Piezo2 acts primarily in F-oDSGCs to segregate the horizontal and vertical OKR pathways.

### Both the loss and overactivation of Piezo2 leads to cross-coupling of the OKR

To determine whether increased Piezo2 activity disrupts AOS circuit function, we next tested the consequences resulting from a gain-of-function (GOF) mutation that prolongs channel opening and delays channel closure, thereby enhancing Piezo2-mediated mechanotransduction (Figure 5A). This *Piezo2^GOF^* conditional mouse line mimics the human *PIEZO2* mutation resulting in distal arthrogryposis 5 (DA5)^[26]^. We generated retina-specific *Piezo2^GOF^* mutant mice (*Ret^Piezo2 GOF^*) using the same *Chx10^Cre^* driver used for our LOF experiments, enabling expression of the mutant channel in retinal progenitor cells. Similarly to *Ret^Piezo2 LOF^* animals, *Ret^Piezo2 GOF^* mice exhibited abnormal cross-coupling in response to temporal motion (Figures 5B–5G), while displaying otherwise normal OKR responses in all directions (Figures 5H–5K). Again, during horizontal ETMs evoked by temporally-moving stimuli, *Ret^Piezo2 GOF^* mice displayed a pronounced vertical component of eye movement, resulting in diagonal trajectories rather than the predominantly horizontal movements observed in control mice (Figures 5B, 5C, 5F and 5G). Notably, these aberrant vertical ETMs, though small, are completely in phase with the larger horizontal ETMs. In contrast, vertical ETMs during nasal stimulation were not significantly altered (Figures 5D and 5E). Together, these findings demonstrate that both insufficient and excessive Piezo2 activity in the retina disrupts the fidelity of horizontal motion processing in the AOS, and they suggest that precise regulation of Piezo2 channel activity is required to maintain the segregation of the horizontal and vertical OKR.

**Figure 5.**
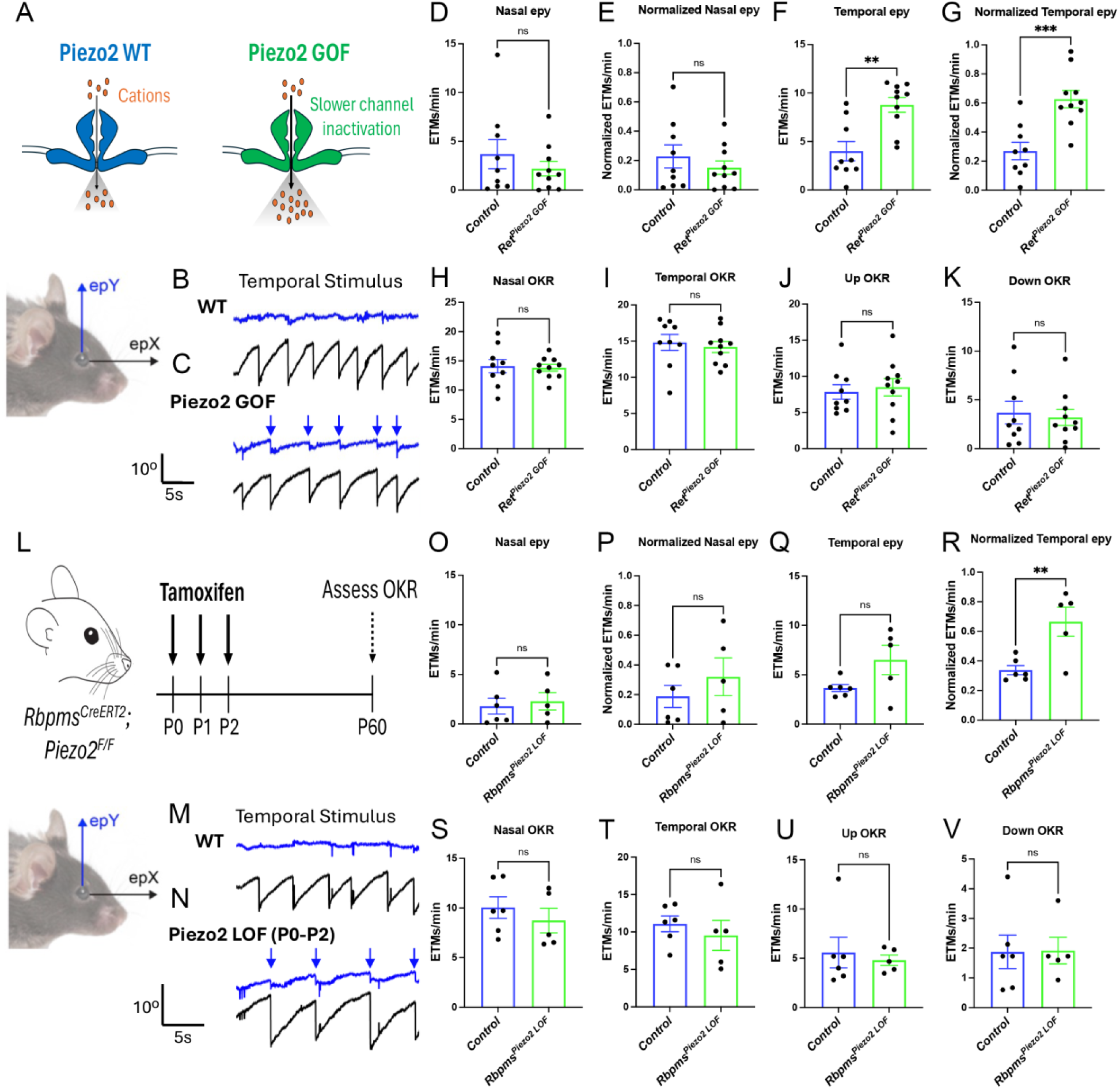
Both loss and overactivation of Piezo2 leads to cross-coupling of the OKR. (**A**) The *Piezo2^GOF^*conditional mouse line harbors a gain-of-function (GOF) mutation that prolongs channel opening and delays channel closure, thereby enhancing Piezo2-mediated mechanotransduction (**B, C**) Retina-specific *Piezo2^GOF^* mice (*Ret^Piezo2 GOF^*) exhibited abnormal cross-coupling in response to temporal stimuli, displaying a pronounced vertical component of eye movement (blue arrows in **C**), indicating that the eye moves diagonally, similar to *Ret^Piezo2 LOF^* mice. (**D, E**) Absolute and normalized number of in-phase upward vertical ETMs/min (nasal epY) during nasal stimuli remain unchanged. (**F, G**) Absolute and normalized number of in-phase upward vertical ETMs/min (temporal epY) during temporal stimuli; *Ret^Piezo2GOF^* mice make significantly more in-phase upward ETMs in response to temporally-moving stimuli than WT mice. (**H-K**) Quantification of ETMs per minute in response to continuous gratings moving in the nasal (**H**), temporal (**I**), upward (**J**), or downward (**K**) direction shows otherwise normal OKR responses in *Ret^Piezo2GOF^* mice. (**L**) Timeline of tamoxifen injections into postnatal mouse pups; control and *Rbpms^CreERT2^, Piezo2^Flox^* pups were injected with tamoxifen from P0-P2 and OKR was assessed at P60. (**M, N**) RGC-specific *Piezo2^LOF^* mice (*Rbpms^Piezo2LOF^*) recapitulate abnormal cross-coupling, with temporal stimuli again eliciting a pronounced vertical component (blue arrows in **N**). (**O, P**) Nasal epY ETMs/min were comparable between *Rbpms^Piezo2LOF^* and control mice in both absolute and normalized measurements. (**Q, R**) During temporal stimuli, *Rbpms^Piezo2LOF^* mice generated significantly more in-phase upward ETMs/min (temporal epY) than control mice in normalized measurements. (**S-V**) Overall OKR responses to continuous gratings moving in the nasal (**S**), temporal (**T**), upward (**U**), or downward (**V**) direction were otherwise normal. Data are presented as mean ± SE. Significance was assessed using Student’s *t*-test. **p < 0.01, ***p < 0.001.

### Piezo2 functions postnatally in retinal ganglion cells to segregate the horizontal and vertical OKR

To define the developmental time window during which Piezo2 acts to establish OKR axis segregation, we deleted *Piezo2* selectively from RGCs during the early postnatal period using an inducible *Rbpms^CreERT2^* mouse line^[27]^, administering tamoxifen at P0-P2 and assessing OKR at P60 (Figure 5L). RGC-specific deletion of Piezo2 during this early postnatal window was sufficient to recapitulate cross-coupling in response to temporally-moving stimuli (Figures 5M–5R) while otherwise normal OKR responses were maintained during continuous checkerboard rotation (Figures 5S–5V). These results indicate that Piezo2 acts cell-autonomously within RGCs, and suggests that it does so during a discrete period of postnatal AOS circuit assembly and refinement.

### Piezo2 regulates developmental refinement of Forward oDSGC projections in the AOS

The segregation of horizontal and vertical OKR pathways is maintained by the precise targeting of RGC axons to distinct AOS nuclei. RGCs encoding horizontal motion project primarily to the NOT, whereas vertical motion-sensitive RGCs project to the MTN. In addition to this anatomical segregation, cross-inhibitory interactions between these two AOS nuclei reinforce the functional separation of horizontal and vertical OKR pathways. Given that loss of Piezo2 results in cross-coupling between horizontal and vertical eye movements, we asked whether this phenotype arises from altered organization of retinal inputs within the AOS.

We first examined whether retina-specific *Piezo2* LOF causes gross changes in retinal projections to retinorecipient brain regions. To visualize RGC axons, we performed intraocular injections of CTB in control and *Ret^Piezo2 LOF^* mice. Analysis of labeled axons revealed robust retinal innervation of the NOT and MTN in both genotypes, with no obvious differences in the density of axonal innervation in any retinorecipient region (Figure 6A). These findings suggest that the cross-coupling phenotype does not arise from large-scale mistargeting of retinal projections to AOS nuclei. We next assessed whether Piezo2 loss affects the morphology of F-oDSGCs. To selectively label NOT-projecting RGCs, we injected a glycoprotein-deleted rabies virus into the NOT and examined retrogradely labeled cells in retinal wholemounts. F-oDSGCs in *Ret^Piezo2 LOF^* mice exhibited dendritic field areas comparable to those observed in control animals (Figures 6B–6D), suggesting that the dendritic architecture of the horizontally-tuned F-oDSGCs remains intact in the absence of Piezo2.

**Figure 6.**
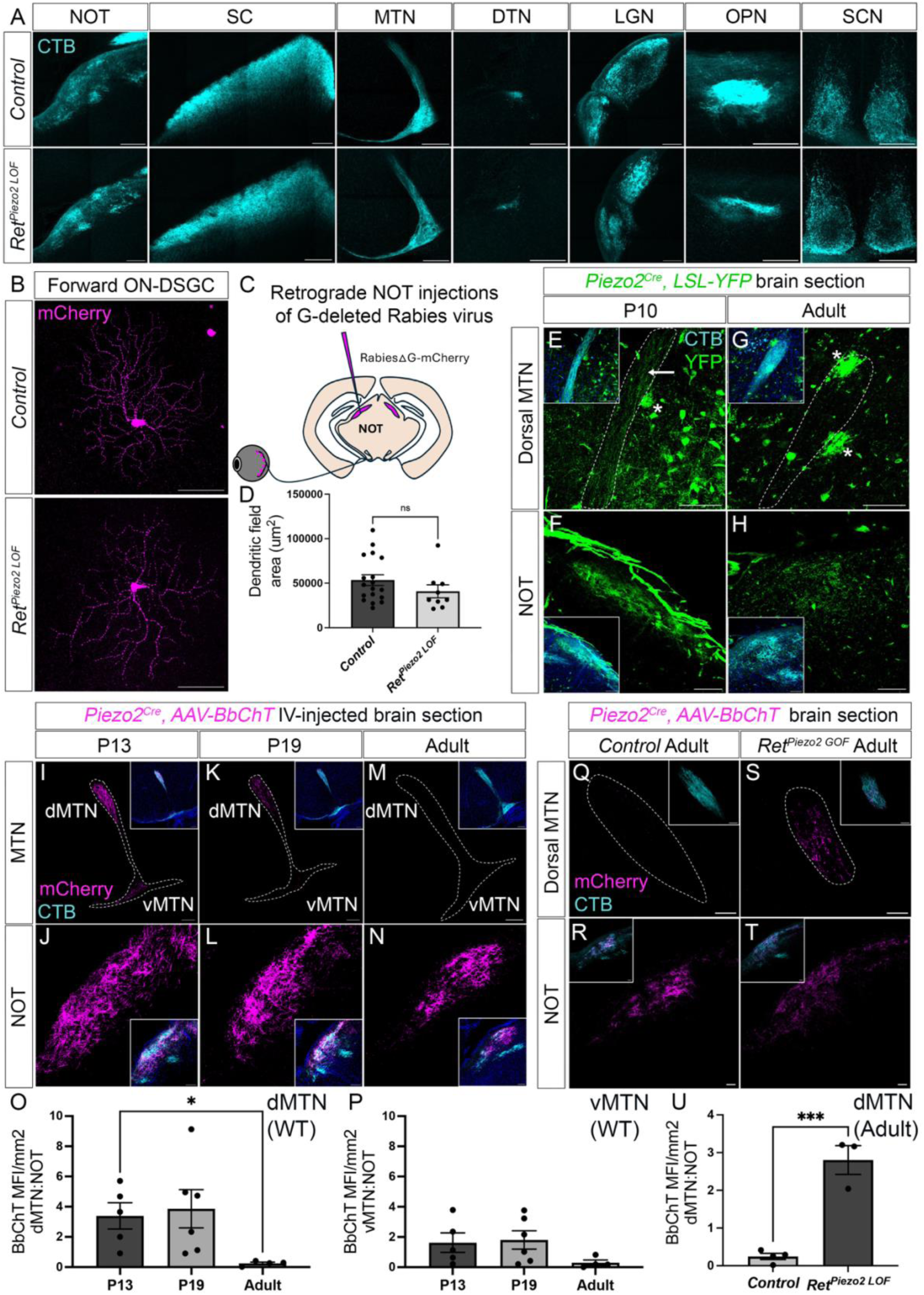
Piezo2 functions in refining axonal projections to the AOS. **(A**) Labeling of retinorecipient targets following intraocular CTB-647 injections in adult WT and *Ret^Piezo2LOF^* mice shows no change in overall innervation. (**B-D**) Adult wholemount retinas depicting viral retrograde filling of NOT-projecting F-oDSGCs; *Ret^Piezo2LOF^* mice show dendritic field areas of F-oDSGCs comparable to those of WT mice. (**E-H**) Intraocular injection of CTB-647 at P2 labels retinorecipient targets at P10, revealing axonal innervation of both the dMTN (**E**) and the NOT (**F**) in *Piezo2^Cre^; LSL-YFP* mice (white arrow). In adults, NOT innervation persists (**H**) while dMTN innervation becomes undetectable (**G**); additional large *Piezo2*-expressing cells visible in the brain are marked with white asterisks and reflect endogenous genetic labeling. (**I-N**) Following intraocular injection of a Cre-dependent AAV encoding mCherry (AAV-BbChT) at P2, Piezo2^+^ RGC axons innervate both the NOT and dMTN at P13 (**I, J**) and P19 (**K, L**) in *Piezo2^Cre^* mice, but dMTN innervation is absent in adult (**M, N**). (**O, P**) Quantification of mean fluorescence intensity of mCherry signal in the dMTN (**O**) and vMTN (**P**), normalized to the NOT of the same animal, across the three timepoints. (**Q-T**) Following intraocular injection of a Cre-dependent AAV encoding mCherry (AAV-BbChT) at P2, Piezo2^+^ axons innervate the NOT in both adult control (**R**) and *Piezo2^Cre/GOF^* (**T**) mice but the dMTN innervation is retained only in *Piezo2^Cre/GOF^* mice (**S**), consistent with a failure of normal axon pruning. (**U**) Quantification of mean fluorescence intensity of mCherry signal in the dMTN, normalized to the NOT of the same animal, reveals persistent dMTN innervation by Piezo2^+^ RGC axons in *Piezo2^Cre/GOF^* mice compared to controls. Data are presented as mean ± SE. Significance was assessed using Student’s *t*-test. *p < 0.05, ***p < 0.001.

Although these results argue against major structural defects in the mature horizontal pathway, the cross-coupling phenotype could arise from more subtle alterations in axonal refinement during AOS circuit development. To explore this possibility, we first examined the normal developmental pattern of Piezo2^+^ RGC projections using *Piezo2^Cre^; LSL-YFP* mice. Analysis of AOS nuclei during the early postnatal period revealed transient Piezo2^+^ innervation of the dorsal MTN. While Piezo2^+^ axons were absent from the MTN in adult mice, we detected YFP-labeled projections in the dorsal MTN at P10 (Figures 6E–6H), suggesting that Piezo2^+^ RGCs initially innervate both horizontal and vertical AOS nuclei before undergoing developmental refinement. Because endogenous *Piezo2* expression throughout the brain complicates visualization of retinal axons in genetically labeled mice, we used a viral labeling strategy to more clearly trace Piezo2^+^ RGC projections. We performed intraocular injections of *AAV-BbChT* in *Piezo2^Cre^* mice at P2 and examined labeled axons at multiple developmental timepoints. Piezo2^+^ axons were observed in the dorsal MTN at P13 and P19, whereas no signal was detected in the MTN of adult mice at P60 (Figures 6I–6N). To control for potential variability in injection efficiency, we normalized the mCherry signal measured in the dorsal MTN to that observed in the NOT for each animal (Figures 6S and 6T). These analyses confirmed a transient developmental innervation of the dorsal MTN by Piezo2^+^ RGCs that is subsequently eliminated during the late postnatal period after eye opening.

One possible mechanism underlying OKR cross-coupling is a disruption of this developmental axonal refinement process. Although the *Piezo2^Cre^* allele does not disrupt Piezo2 function and therefore cannot be used to assess Piezo2-specific projections under LOF conditions, we were able to examine the effect of increased Piezo2 activity using mice carrying one copy of the *Piezo2^GOF^* allele driven by *Piezo2^Cre^*. Therefore, we performed intraocular injections of *AAV-BbChT* in *Piezo2^Cre/GOF^* mice at P2 and examined their axonal projections at adulthood (P60). Strikingly, we observed the persistence of Piezo2^+^ axons in the dorsal MTN of *Piezo2^GOF/+^* mice, indicating that excessive Piezo2 activity interferes with the normal developmental elimination of these projections (Figures 6Q–6U).

Together, these results suggest that Piezo2 channel activity contributes to the developmental refinement of F-DSGC projections within the AOS.

## DISCUSSION

Here we report a vital role for Piezo2 in the behavioral segregation of horizontal and vertical OKR responses. We characterized *Piezo2* expression in AOS-projecting F-DSGCs that include F-oDSGCs and the recently described perivascular RGCs^[23]^, which we identify as the slow F-ooDSGCs first characterized over a decade ago^[3]^. RGC-specific removal of *Piezo2* produces a cross-coupled OKR phenotype in which ectopic upward ETMs are elicited by horizontally-moving stimuli. We show that Piezo2 function in slow F-ooDSGCs is entirely dispensable for preventing OKR cross coupling, suggesting that Piezo2 acts in F-oDSGCs to segregate the horizontal and vertical OKR. Strikingly, both loss and gain of Piezo2 function drive this cross-coupled OKR, revealing that precise titration of Piezo2 channel activity appears critical for correct AOS circuit function. To account for this phenotype, we document early postnatal misprojections from *Piezo2*-expressing RGCs to the MTN that are normally pruned, by adulthood, and show that these ectopic projections persist when Piezo2 activity is elevated in the retina. Such aberrant co-activation of horizontal and vertical pathways provides a plausible anatomical mechanism for the cross-coupling phenotype observed in *Piezo2* mutant mice (Figure 7). Together, these findings support a model in which aberrant MTN activation by horizontal motion, due to failed pruning of ectopic projections, drives the inappropriate recruitment of vertical eye movements during the horizontal OKR (Figure 7).

**Figure 7.**
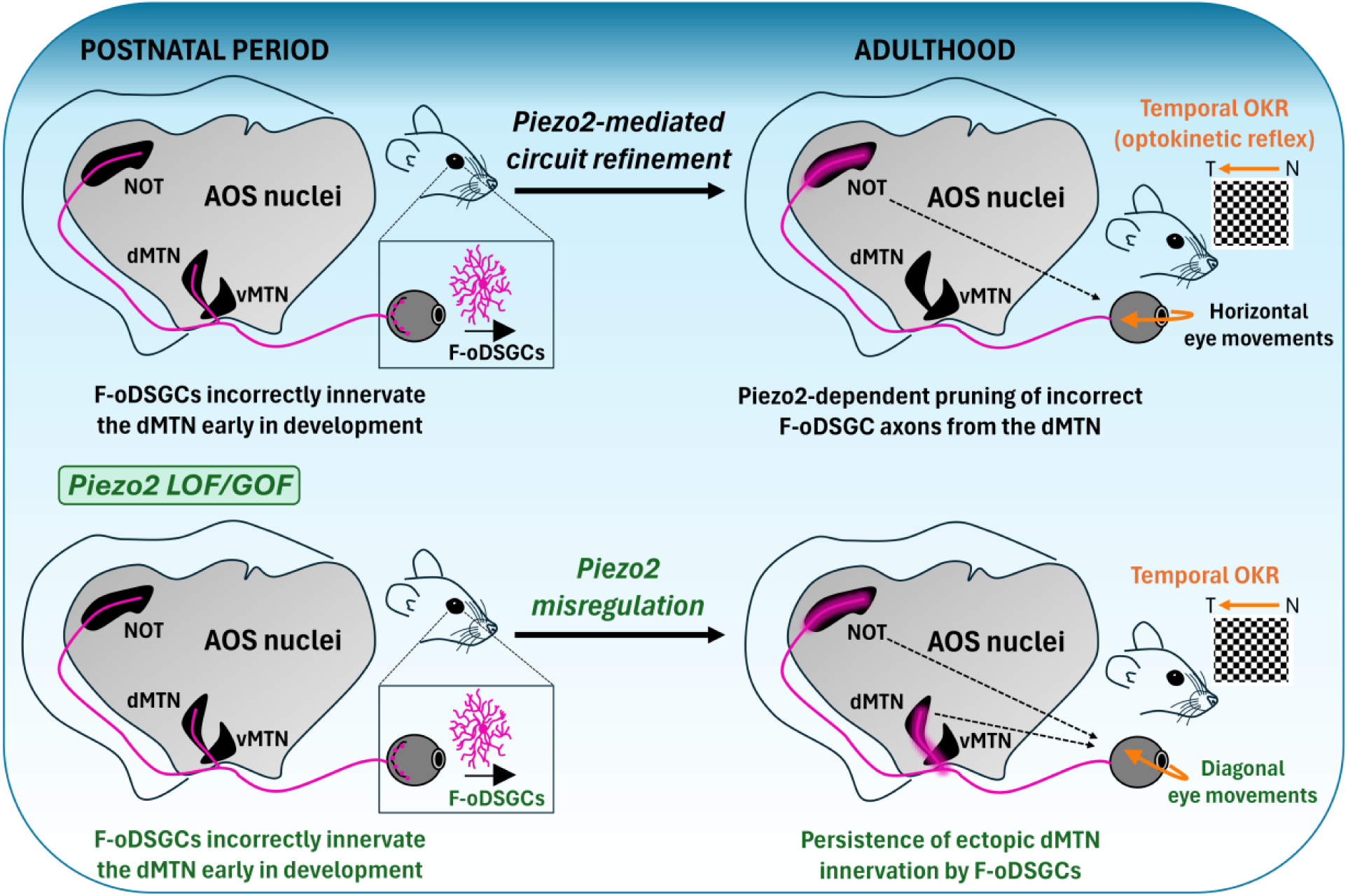
Piezo2-mediated axon pruning in F-oDSGCs is required for precise AOS circuit function (Top) During the postnatal period, F-oDSGC axons transiently innervate both the NOT (the F-oDSGC target nucleus) and the dMTN (the U-oDSGC target nucleus). Piezo2-dependent circuit refinement subsequently prunes ectopic F-oDSGC projections from the dMTN, resulting in adult F-oDSGC axons that innervate the NOT (and the DTN, not depicted) exclusively. This precise targeting is required for normal horizontal OKR responses to temporally moving stimuli. (**Bottom**) In Piezo2 LOF and GOF mice, misregulation of Piezo2 channel activity disrupts axon pruning, resulting in the persistence of ectopic F-oDSGC innervation of the dMTN into adulthood. This aberrant connectivity drives simultaneous activation of both the NOT and the dMTN by F-oDSGCs in response to horizontally moving stimuli, cross-coupling the horizontal and vertical OKR and manifesting as diagonal eye movements in response to temporal stimuli.

### Developmental refinement of the AOS

The assembly of visual circuits requires a series of events including initial axon targeting, precise synaptic partnering, and in many cases the subsequent pruning and refinement of projections^[28, 29]^. In the AOS circuit, oDSGCs start to innervate their target nuclei during late embryogenesis, and this innervation continues postnatally^[13, 14]^. Some molecular pathways directing the assembly of AOS circuits have been identified. Sema6a/PlexA2/A4 signaling directs vertically-tuned oDSGCs to the MTN and also is important for interconnectivity between the MTN and NOT^[14, 30]^, while horizontally-tuned F-DSGCs are guided to the NOT through Cntn4/APP interactions^[15]^.

Despite progress in understanding initial targeting, evidence for subsequent refinement within the AOS has remained sparse. Prior work documents to some degree postnatal pruning in AOS nuclei between P8 and P20, but these ectopic projections arise from fast-tuned nasal ooDSGCs, a population not associated with AOS function, which transiently innervate the NOT^[15]^. Previous work describes early-born RGC subtypes whose axons overshoot their targets and undergo removal prior to P20 via a combination of cell death and axon retraction, and in this study, it was found that AOS-projecting RGCs do not innervate targets outside of the AOS^[31]^. Importantly, however, this study used *Hoxd10-GFP* to label all AOS-projecting RGCs and therefore could not resolve whether, within AOS nuclei, individual oDSGC subtypes transiently mistarget to the wrong AOS nucleus. The emergence of subtype-specific markers and genetic tools enabling access to distinct oDSGC populations has only recently made such assessments possible. Here, leveraging these advances, we report that slow horizontally-tuned F-DSGC axons transiently mistarget to the MTN, the vertical-axis AOS nucleus, during the early postnatal period. This ectopic innervation is present at P13 and P19, but is absent in adults, demonstrating that the AOS does undergo subtype-specific circuit refinement previously undetectable at the resolution of bulk DSGC labeling. Crucially, the failure of this refinement, manifested as the persistence of only a few ectopic F-DSGC axons in the MTN, is sufficient to produce cross-coupling of the horizontal and vertical OKR.

What signals drive this postnatal AOS refinement? One possibility is that refinement is instructed directly by the onset of visual experience. Eye opening in mice occurs around P13, coinciding with the temporal window during which we observe ectopic MTN innervation. Indeed, visual experience-dependent refinement of RGC axons is well documented in other retinorecipient targets. Development of retinogeniculate connections passes through distinct refinement phases, including a prolonged phase of visual experience-dependent refinement that extends well into the fourth postnatal week and beyond^[32]^. Taking this into account, patterned sensory input itself could provide the instructive signal that stabilizes appropriate projections while eliminating inappropriate ones. Testing this model by dark-rearing animals through the refinement window represents an important future direction. Another possibility is that refinement is driven not simply by sensory exposure, but by correlated activity generated during the emergence of functional circuits and their associated behaviors. Across cortical and subcortical brain regions, correlated neuronal activity can instruct axonal and dendritic refinement through Hebbian-like competitive mechanisms, and the developing visual system is no exception^[17, 28, 33, 34]^. Within the AOS, the onset of the OKR after eye opening could result in correlated firing among U-oDSGCs converging onto the dorsal MTN during vertical image motion. Such coordinated activity may selectively reinforce canonical vertical-motion inputs, rendering the ectopic F-DSGC inputs out of sync and thus triggering their selective elimination.

### A mechanosensory brake on ectopic axonal connectivity

Piezo2 has been implicated in gentle touch, proprioception^[22, 35, 36]^ and mechanosensory processes in a wide range of internal organs, including the lung^[37]^, aorta^[38, 39]^, bladder^[40]^, kidney^[41]^, and adipose tissue^[42]^. Beyond these canonical mechanosensory roles, Piezo channels have been implicated in neural circuit development and regeneration. Knockdown of *piezo1* impairs RGC targeting to the optic tectum in *Xenopus* embryos^[43]^, while loss of *DmPiezo* enhances axon regeneration of *Drosophila* class III dendritic arborization neurons and mouse corneal sensory neurons^[44]^, suggesting that Piezo channels can both guide initial axon targeting and later suppress regrowth. Piezo channels also regulate various aspects of dendritic morphogenesis and outgrowth. Overexpression of the *C. elegans* homolog *pezo-1* rescues impaired Ca^2+^ transients and dendritic elaboration in DEG/ENaC mutant PVD neurons through an intact pore domain, implicating ion channel activity in the transduction of mechanical forces during dendrite outgrowth^[45]^. Intriguingly, Piezo channels can also regulate neuronal wiring independently of mechanosensory activity. *DmPiezo* was identified in a proteomic screen as a cell surface protein regulated by the transcription factor *Acj6* and required for dendritic targeting of olfactory projection neurons^[46]^. A pore-dead variant fully rescued the targeting deficits of *DmPiezo* mutants, demonstrating that channel activity is dispensable for this wiring function. Together, these studies strongly implicate Piezo2 in neuronal circuit wiring and provide a rich precedent for the Piezo2 role we describe here in axonal refinement within the AOS.

In most biological contexts, loss and gain of Piezo2 activity produce opposing phenotypes^[42, 47]^; however, in rare cases they have been shown to drive similar outcomes^[39]^. Both loss and gain of Piezo2 activity lead to cardiac hyperplasia and altered coronary artery formation^[39]^. Here, we similarly find that both Piezo2 LOF and overactivation produce a cross-coupled OKR phenotype, raising the question of whether they act via shared or distinct mechanisms. One possibility is that they produce the same behavioral outcome via distinct circuit-level perturbations. For instance, the NOT and the MTN are reciprocally connected, and this interconnectivity normally permits mutual inhibition such that vertical eye movements are suppressed during horizontal OKR and vice-versa^[30]^. Piezo2 LOF might therefore disrupt information flow to the NOT that in turn alters its inhibitory connections with the MTN, independently of the ectopic F-DSGC projections to the MTN we observe upon Piezo2 overactivation. An alternative possibility is that both manipulations converge on the same mechanism to disrupt developmental elimination of F-DSGC projections from the MTN. In this model, a precise level of Piezo2 activity is required to drive the pruning of these transient inputs, and perturbation in either direction disrupts this process. In either case, the recapitulation of the LOF behavioral phenotype upon Piezo2 overactivation indicates that its role in F-DSGC axonal refinement is dependent on its channel activity, distinguishing it mechanistically from the channel-independent wiring function described in the fly olfactory system^[46]^.

How might Piezo2 activity regulate axon pruning? Two interconnected downstream effectors are particularly relevant. Piezo2 is functionally linked to the actin cytoskeleton via filamin-B, which serves as a molecular tether^[48]^. Piezo2-mediated Ca2^+^ influx has also been shown to activate RhoA to control the formation and orientation of active stress fibers and focal adhesions^[49]^. These cytoskeletal connections position Piezo2 as a direct regulator of the axon terminal mechanical state, which is known to influence branch stability and retraction^[50, 51]^. Second, tightly regulated intracellular Ca2^+^ signaling is a central mediator of developmental axon refinement. Local calcium transients regulate axonal branch dynamics by controlling cytoskeletal remodeling and synaptic stabilization, and both insufficient and excessive calcium activity disrupt pruning processes; Ca2^+^ elevations suppress growth and promote axon retraction^[52]^, while oscillatory Ca2^+^ signals via voltage-gated Ca2^+^ channels are required for the activity-dependent elimination of ectopic contacts during synaptic refinement^[53]^. In this framework, loss of Piezo2 could reduce calcium influx needed to engage pruning pathways that eliminate transient F-DSGC projections to the dorsal MTN, while prolonged channel opening in *Piezo2^GOF^* mutants might produce sustained calcium elevations that paradoxically stabilize immature axonal branches or disrupt the temporal dynamics of calcium signaling required for refinement. Both scenarios would result in persistence of ectopic MTN inputs and the cross-coupled OKR phenotype we observe.

These potential modes of Piezo2 action may further extend to transcriptional regulation. Recent work shows that *Piezo2* deletion in sensory neurons is associated with broad transcriptional changes^[54, 55]^, including altered expression of axon guidance molecules such as Nrp2, EphA4, and Sema5a^[54]^. Notably, our finding that overactivation of Piezo2 produces ectopic MTN innervation and a similar behavioral phenotype as *Piezo2^LOF^* mutants suggests that if transcriptomic changes in axon guidance genes contribute to mistargeting in F-DSGCs, they are likely regulated by the level of Piezo2 activity rather than simply the presence or absence of the channel.

Dissecting the precise mechanisms by which both loss and gain of Piezo2 activity produce OKR cross-coupling will require new genetic tools that provide selective access to F-DSGCs, enabling simultaneous conditional knockout of Piezo2 and visualization of F-DSGC axons.

### Functional divergence among AOS-projecting DSGC subtypes

The existence of two forward-preferring subtypes projecting to the NOT (F-oDSGCs and slow F-ooDSGCs) has been recognized for over a decade^[3]^, yet their relative contributions to the horizontal OKR have remained unresolved. Here we provide both molecular and functional evidence that these populations are not redundant parallel inputs to the NOT but instead serve distinct roles in AOS development.

A key step toward resolving this question was the identification of a molecular marker for slow F-ooDSGCs. A recent study discovered a perivascular RGC subset defined by *Nts* expression and showed that Piezo2 functions in these cells regulate the correct patterning of penetrating blood vessels^[23]^. We show here that these Nts^+^ RGCs likely correspond to the slow F-ooDSGCs described previously, providing a molecular handle for distinguishing this population from horizontal oDSGCs. Using an *Nts^Cre^* mouse line, we find that Nts^+^ RGCs constitute a homogenous population of bistratified ON-OFF RGCs whose dendritic arbors co-stratify with SACs, confirming their identity as ooDSGCs. We further show that Nts^+^ RGCs project to the NOT and that a subpopulation is co-labeled in the *Hoxd10-GFP* mouse line, confirming their identity as AOS-projecting slow F-ooDSGCs.

Despite the previously established requirement for Piezo2 in Nts^+^ RGCs for retinal vascular patterning^[23]^, we find that Piezo2 function in slow F-ooDSGCs is entirely dispensable for normal OKR responses. In addition to slow F-ooDSGCs, *Piezo2* is also expressed in F-oDSGCs and amacrine cells in the retina. We demonstrate here that selective removal of *Piezo2* from amacrine cells fails to recapitulate the cross-coupled OKR phenotype, while removal from RGCs is sufficient to drive it. Together, these results strongly suggest that Piezo2 acts in F-oDSGCs to segregate the horizontal and vertical OKR.

These findings provide a clear functional dissociation between these two forward-preferring DSGC populations. While both subtypes project to the AOS and express *Piezo2*, they use Piezo2 to regulate mechanistically distinct functions, suggesting that they serve different developmental functions and likely encode different aspects of the horizontal OKR. F-oDSGCs are canonical slow-speed tuned oDSGCs optimally suited to drive the smooth tracking phase of the OKR, while slow F-ooDSGCs are capable of responding across a broader velocity range^[3]^ and may contribute to the horizontal OKR under distinct stimulus conditions. Complete resolution of their respective functional contributions requires the generation of genetic tools providing selective access to F-oDSGCs, enabling the selective ablation of each population to directly characterize their distinct contributions to the horizontal OKR.

### Implications for eye movement disorders

Several human diseases caused by mutations at the *PIEZO2* locus underscore its developmental importance across multiple organ systems. Gain-of-function mutations that slow channel inactivation and cause hyperactivity, including the human variant represented by the humanized *Piezo2^GOF^*mouse line used here^[26]^, underlie distal arthrogryposis type 5 (DA5), an autosomal dominant disorder featuring joint contractures and restrictive lung disease^[56]^. Overlapping phenotypes are seen in Gordon syndrome and Marden-Walker syndrome, which are also caused by dominant *PIEZO2* mutations and likely represent phenotypic variants of the same condition^[57]^. Importantly, individuals with *PIEZO2* variants have been reported to have ocular motor phenotypes including ophthalmoplegia, restrictions in horizontal eye movements, and restricted upward gaze^[58]^. The mechanistic basis of these oculomotor deficits has historically been attributed to extraocular muscle dysfunction or abnormal cranial nerve innervation^[59, 60]^. Our findings raise the alternative, or complementary, possibility that disrupted Piezo2 function in retinal neurons contributes to the eye movement phenotypes by impairing AOS circuit assembly. Indeed, *Piezo2* expression in RGC subsets, including the perivascular RGCs we identify here as slow F-ooDSGCs, is conserved from mouse to human^[23]^, suggesting possible functional conservation as well. Careful recordings of the human optokinetic nystagmus (OKN) in individuals with *PIEZO2* variants will be an important next step in determining whether Piezo2 plays a conserved role in the segregation of the horizontal and vertical OKR, and whether aberrant cross-coupling of these axes contributes to the eye movement phenotypes seen in PIEZO2-related diseases.

## ACKNOWLEDGEMENTS

We are grateful to Andrew Huberman, Masaharu Noda, Seth Blackshaw, Xin Duan for generously providing *Hoxd10-GFP, Spig1-GFP, Chx10^Cre^, and Nts^Cre^* mice, respectively. We thank all members of the Kolodkin laboratory for comments and helpful discussions. This work was supported by R01 EY032095 (ALK) and Research to Prevent Blindness (RPB) Stein Innovation Award.

## LEAD CONTACT

Further information and requests for resources and reagents should be directed to and will be fulfilled by the lead contact, Alex L. Kolodkin.

## METHODS

### Experimental model and subject details

All animals were approved by the Institutional Animal Care and Use Committee (IACUC) at the Johns Hopkins University School of Medicine. The day of birth was designated as postnatal day 0 (P0). Mice of both sexes were used for experiments. *Hoxd10-GFP* mice were a gift from A. Huberman and were described previously^[3]^. *Spig1-GFP* mice were a gift from M. Noda and were described previously^[12, 13]^. *Chx10^Cre^* mice were a gift from S. Blackshaw and were described previously^[24]^. *Nts^Cre^* mice were a gift from X. Duan and were described previously^[62]^. *Rbpms^CreERT2^* mice were obtained from L. Gan and were described previously^[27]^. *Piezo2^GOF^* mice were obtained from A. Patapoutian and were described previously^[26]^. *Piezo2^Cre^* (Strain #027719)*, Piezo2^Flox^* (Strain #027720)*, Ptf1a^Cre^* (Strain #023329)*, Ai14* (Strain #007914)*, Ai2* (Strain #007920), and *MORF3* (Strain #035403) mice were obtained from Jackson Laboratories. All mice were maintained on a C57BL/6J background. Animals were housed in a 12-hour light-dark cycle and visual behavioral tests were performed during the light cycle.

### Intraocular CTB and AAV injections

For intraocular injections of adult mice, mice were anesthetized with 2% isoflurane and secured in a stereotaxic apparatus. A shar 30G needle was used to make a hole at the corneal limbus and 1μl of either AAV, CTB, or a 1:1 combination of both, was injected into the vitreous using a pulled glass pipette. The following reagents were used: AAV9-BbChT (Addgene, stock #45186-AAV9), AAV2 CAG-FLEX-GFP (UNC Vector Core), cholera toxin subunit B Alexa Fluor 555 conjugate (1 mg/mL, ThermoFisher Scientific), and cholera toxin subunit B Alexa Fluor 647 conjugate (1 mg/mL, ThermoFisher Scientific). For intravitreal injections of early postnatal (P1-P3) mice, mice were anaesthetized on ice for 4 minutes. An incision was made using a sharp 30 G needle to expose the eyeball. A microinjector set to a pressure of 20 psi and a 35 ms pulse duration was used to deliver nanoliter volumes of AAV, CTB, or both into each eye. Mice injected with AAV were sacrificed after 2 weeks, and mice injected with CTB were sacrificed after at least 2 days.

### Immunohistochemistry

To stain whole-mount retinas, mice were perfused with phosphate buffered saline (PBS) followed by 4% paraformaldehyde (PFA). Eyeballs were post-fixed in 4% PFA for 1 hour at room temperature, or overnight at 4°C, and then washed at least 3 times with PBS. Retinas were dissected in PBS and incubated for 4 hours at room temperature in permeabilization buffer containing 3% bovine serum albumin, 0.3% Triton X-100, and 0.02% sodium azide. Following permeabilization, retinas were rinsed for 3 hours in PBS-T (PBS, 0.4% Triton X-100). Antibodies were diluted in blocking solution containing 20% DMSO and 10% donkey or goat serum in PBS-T. Retinas were incubated with primary antibodies for 3-5 days at 4°C, washed 6 times with PBS-T, and incubated with secondary antibodies for 2 days at 4°C. Finally, stained retinas were washed 5 times with PBS-T and flat-mounted on Superfrost microscope slides.

To stain retinal cross sections, mice were perfused with PBS followed by 4% PFA, and enucleated eyeballs were post-fixed in 4% PFA for 1 hour at room temperature, or overnight at 4°C. After washing the eyeballs 3 times with PBS, an incision was made on the cornea using a scalpel. Eyeballs were incubated in a sucrose series with 10%, 20%, and 30% sucrose in PBS at 4°C until sinking. Cryopreserved eyeballs were then embedded and frozen in Neg-50 embedding medium (Richard-Allen Scientific, Kalamazoo, MI). A cryostat was used to make retinal cross sections of 25 μm thickness. Prior to staining, slides were dried for 30 minutes at room temperature and washed for 5 minutes in PBS. Cross-sections were blocked for 1 hour at room temperature in blocking buffer consisting of 10% donkey or goat serum and 0.1% Triton X in PBS. Antibodies were diluted in a dilution buffer composed of 1% donkey or goat serum and 0.1% Triton X in PBS. In a humidified chamber, retinal cross-sections were incubated with primary antibodies overnight at 4°C, washed 3 times with PBS, and incubated with secondary antibodies for 2 hours at room temperature. Stained cross sections were washed 3 times with PBS prior to mounting.

To stain brain sections, mice were perfused with PBS followed by 4% PFA. Brains were removed and post-fixed in 4% PFA overnight at 4°C, then washed 3 times with PBS. For vibratome sectioning, brains were embedded in a 3% (w/v) solution of low melting point agarose in PBS. Brains were sliced coronally at a thickness of 200 μm and the sections were then incubated in permeabilization buffer, consisting of 3% bovine serum albumin, 0.3% Triton X-100, and 0.02% sodium azide in PBS, overnight at 4°C. Permeabilized brain sections were incubated with primary antibodies diluted in permeabilization buffer overnight at 4°C, washed 8 times with PBS, and incubated with secondary antibodies overnight at 4°C. Before mounting, brain sections were washed 3 times in PBS. All images were acquired with a Zeiss LSM 700 confocal microscope.

Antibodies used for immunohistochemistry include: chicken anti-GFP (AVES, 1:1000), rabbit anti-DsRed (Living Colors, 1:1000), rabbit anti-V5 (Bethyl Laboratories, 1:400), guinea pig anti-RBPMS (PhosphoSolutions, 1:500), guinea pig anti-RBPMS (ThermoFisher, 1:500), rabbit anti-RBPMS (Proteintech, 1:500), goat anti-ChAT (Millipore, 1:200), goat anti-VAChT (Millipore [ABN100], 1:200), guinea pig anti-VAChT (Synaptic Systems, 1:500), rabbit anti-Calbindin (Swant [CB38], 1:2000), rabbit anti-Calretinin (Swant [CR7697], 1:1250), mouse anti-Goα (Chemicon, 1:500), and guinea pig anti-vGlut3 (Synaptic Systems, 1:1250).

### Fluorescence *in situ* hybridization

RNAscope *in situ* hybridization of fixed retinal tissue was performed according to manufacturer’s instructions with slight modifications (Advanced Cell Diagnostics, Newark, CA). Juvenile mice were anesthetized with a 2.5% working solution of avertin and perfused with RNase-free PBS followed by 4% PFA. Enucleated eyes were post-fixed in 4% PFA overnight at 4°C, then washed three times with RNase-free PBS. A scalpel was used to cut a slit in the cornea, and eyes were cryopreserved in series with 10%, 20%, and 30% (w/v) RNase-free sucrose/PBS. Eyes were then frozen in Neg-50 embedding medium (Richard-Allen Scientific, Kalamazoo, MI), and sectioned at a thickness of 14 μm. To preserve the fluorescence of endogenous fluorophores, slides were baked for 30 minutes at 45oC, rather than at 60oC, prior to performing the RNAscope assay. Probes targeting *Ptprt*, *Ptprk*, *Fibcd1*, and *Rbpms* were provided by the manufacturer. Positive RNAscope signal appears as punctate dots. Dotted lines were drawn around the collection of mRNA puncta for the gene of interest. All images were acquired with a Zeiss LSM 700 confocal microscope.

### Headpost installation surgery

Headpost surgery was performed on mice older than 8 weeks of age. Mice were anesthetized with 2% isoflurane and positioned in a stereotaxic apparatus. Four burr holes were drilled into the skull on either side of the sagittal suture between bregma and lambda to allow for the placement of four 1.00UNM ’ 0.120” stainless steel screws. A pedestal, serving as the base for an acrylic headpost, was made by placing dental acrylic around the four screws (Ortho-Jet, Lang Dental; Wheeling, Illinois, USA). Headposts were fabricated from custom molds incorporating two M1.4 hex nuts embedded in dental cement. During surgery, the headpost was placed over the acrylic pedestal. Mice were allowed to recover for at least 7 days before visual behavior assessments.

### Optokinetic reflex recording

The OKR recording chamber consists of a square box containing four computer monitors, each 12 inches tall by 20 inches wide, that project a black-and-white checkboard with a stripe width subtending 5° of visual angle. While mice were restrained and head-fixed in a custom holder, an infrared video camera was used to record eye movements. Calibration was performed for each mouse to transform the linear displacement of the pupil in millimeters to rotational displacement in degrees^[63]^. Continuous and sinusoidal stimuli were presented in both horizontal and vertical directions. Each trial for continuous stimuli consisted of 10 cycles, alternating between a checkboard rotating at 5°/s for 30s, and a grey screen presented for 30s. Sinusoidal stimuli were presented at a temporal frequency of 0.2 Hz and amplitude of 5° of visual angle, for a total of 10 cycles per trial. OKR recordings were analyzed using Igor Pro v6 (WaveMetrics, Inc., Portland, OR). Eye tracking movements (ETMs), defined as one slow phase followed by one saccade^[9, 19]^, were manually counted and used to calculate the rate of ETMs (ETMs/minute) for continuous stimuli. Cross-coupled ETMs were identified as those saccades in the vertical trace that were completely in phase with the larger saccades in the horizontal trace. The number of these coinciding ETMs were then normalized to the total number of horizontal ETMs in that trace. For sinusoidal stimuli, saccades were manually removed, and the gain of eye movement was calculated using a custom script^[1]^. Gain is defined as the ratio of the angular velocity of the pupil to the angular velocity of the stimulus.

### Looming assay

Mice were dark-adapted for 1 hour prior to being placed in the looming chamber. The looming chamber consists of a transparent plexiglass box (10 inches wide x 20 inches long x 15 inches tall) with an overhead computer monitor. The floor of the looming chamber contains a triangular prism-shaped shelter (4.5 inches per side by 6.25 inches long) on the far side and a marked square (4.5 inches x 4.5 inches) at the center. Looming assays were performed in a dark room with light emanating only from the overhead monitor of the looming chamber. Each mouse was acclimated to the looming chamber for 10 minutes prior to the presentation of the looming stimulus. After the acclimation period, if the mouse contacted the marked square at the center of the floor, a looming stimulus was presented on the overhead monitor. The looming stimulus consists of an expanding black circle on a white background, appearing 15 times over a period of 10 seconds. A video camera with infrared/night vision capability was used to record the looming response. A normal looming response was defined as seeking shelter or freezing. The latency to response in seconds was recorded for each mouse.

### Stereotactic surgery and NOT injections

Adult mice were anesthetized using 2% isoflurane and placed into a stereotactic apparatus. Rabies virus SAD-B19-RVdG-tdTomato (3.50E+10; UC Irvine Vector Core, Cat # RVdG-5) was injected into the nucleus of the optic tract (NOT) (150 nl at 10 nl/s) using a Hamilton Neuros syringe coupled to a computerized microsyringe pump controller (World Precision Instruments). Five minutes elapsed between the injection and removal of the needle to allow diffusion of CTB around the injection site. The NOT coordinates are as follows: anterior/posterior -2.6 mm, medial/lateral ±1.0 mm, dorsal/ventral -2.2 mm (from surface), (coordinates are relative to bregma). Following injection, five minutes were allowed to elapse before removing the syringe to allow for diffusion to occur. Rabies-injected mice were housed for 5 days to allow adequate transsynaptic spread.

### Single-cell RNA sequencing cell isolation

For single-cell RNA sequencing of *Hoxd10-GFP* retinal ganglion cells, GFP+ cells were isolated by fluorescence-activated cell sorting (FACS) and RGCs were positively selected using a CD90.2-BV421 antibody. In total, 1,160 single cells were sequenced across five developmental time points, from embryonic day 18.5 (E18.5) to postnatal day 21 (P21), at an average depth of 1.1 × 10⁶ ± 0.2 × 10⁶ reads per cell, with 90 ± 2% alignment to the genome. Single-cell cDNA libraries were generated using the Smart-seq2 protocol^[64]^ and sequenced on an Illumina NextSeq 500 platform with 75-bp paired-end reads.

### Quantification and statistical analysis

Graphs were generated using the ggplot2 package in R v3.5.1 (The R Foundation for Statistical Computing, Auckland, New Zealand) and GraphPad Prism 9.5.0. Reanalysis of a P5 RGC transcriptomic analysis was done as described previously^[61]^. RNA FISH images were quantified using thresholding and automated particle analysis in ImageJ. Particle size parameters were set between 0.04-2, and cells were considered positive if they contained >3 puncta. ImageJ software was used to trace virally labeled RGC dendrites with the simple neurite tracer (SNT) package and to count cells using the Cell Counter plugin. Dendritic field areas were measured by drawing a polygon connecting the dendritic tips of virally labeled RGCs. Axonal innervation of virally labeled Piezo2^+^ RGCs was quantified by first identifying and outlining the innervated brain nucleus (dMTN, vMTN, or NOT) based on CTB labeling. Within this outlined region, fluorescence intensity was measured across Z-stacks and background signal was subtracted. To account for variability in nucleus size across animals, the intensity value was normalized to the area of the outlined region. To further account for variability in viral injection efficiency, the normalized dorsal and ventral MTN values were then normalized to the corresponding NOT value from the same animal. Significance was determined with either a *t*-test or a one-way ANOVA with a post-hoc Tukey’s test unless otherwise specified. Significance was defined as p < 0.05.

### Single-cell RNA sequencing data analysis

Single-cell RNA sequencing FASTQ files were mapped to the GRCm38.p6/mm10 mouse reference genome using HISAT2 v2.1.0^[65]^ and the resulting alignment files were processed with SAMtools v1.3.1^[66]^. Transcript abundance was estimated with Cuffquant and normalized with Cuffnorm, both from Cufflinks v2.2.1^[67]^.

Downstream analyses were performed in R v 3.6.2 using Seurat^[68, 69]^. An annotated expression matrix was generated from Cuffnorm FPKM output, and a Seurat object was created using genes detected in at least three cells and cells with at least 200 detected features. Twenty outlier cells identified during initial processing were removed before downstream analysis. To exclude non-target retinal populations, cells with detectable expression of the rod photoreceptor markers Rcvrn or Pde6g, the ipRGC markers Opn4 or Eomes, the microglial marker Cx3cr1, the amacrine cell marker Prox1, or the bipolar cell markers Otx2 or Vsx2 were removed. Retained cells were required to express the RGC marker Rbpms. Additional quality-control filtering retained cells with 5,000-12,500 detected genes.

Expression values were log-normalized using a scale factor of 10,000, and 2,000 variable features were identified for downstream analysis. Data were scaled, and developmental time point was regressed out before dimensionality reduction and clustering. Principal component analysis was performed using variable features, and the first 20 principal components were used for nearest-neighbor graph construction, clustering, and UMAP visualization. Cells were clustered at a resolution of 0.8. Cluster markers were identified using Seurat’s FindAllMarkers function, retaining positive markers expressed in at least 25% of cells in a cluster with a log-fold-change threshold of 0.25. Cluster identities were assigned based on expression of known DSGC subtype markers, including Ptprk in U-oDSGCs and Ptprm in D-oDSGCs.

## REFERENCES

1. Kodama, T. and S. du Lac, Adaptive Acceleration of Visually Evoked Smooth Eye Movements in Mice. J Neurosci, 2016. 36(25): p. 6836–49.

2. Simpson, J.I., The accessory optic system. Annu Rev Neurosci, 1984. 7: p. 13–41.

3. Dhande, O.S., et al., Genetic dissection of retinal inputs to brainstem nuclei controlling image stabilization. J Neurosci, 2013. 33(45): p. 17797–813.

4. Harris, S.C. and F.A. Dunn, Asymmetric retinal direction tuning predicts optokinetic eye movements across stimulus conditions. Elife, 2023. 12.

5. Hamilton, N.R., A.J. Scasny, and A.L. Kolodkin, Development of the vertebrate retinal direction-selective circuit. Dev Biol, 2021. 477: p. 273–283.

6. Giolli, R.A., R.H. Blanks, and F. Lui, The accessory optic system: basic organization with an update on connectivity, neurochemistry, and function. Prog Brain Res, 2006. 151: p. 407–40.

7. Hoffmann, K.P., Comparative neurobiology of the optokinetic reflex in mammals. Rev Bras Biol, 1996. 56 Su 1 Pt 2: p. 303–14.

8. Kamermans, M., et al., A retinal origin of nystagmus-a perspective. Front Ophthalmol (Lausanne), 2023. 3: p. 1186280.

9. Yonehara, K., et al., Congenital Nystagmus Gene FRMD7 Is Necessary for Establishing a Neuronal Circuit Asymmetry for Direction Selectivity. Neuron, 2016. 89(1): p. 177–93.

10. Qi, J., A. Mastumoto, and K. Yonehara, Distinct optokinetic reflex phenotypes in Frmd7 and Chrnb2 mutant mice. bioRxiv, 2026: p. 2026.04.03.716267.

11. Vaney, D.I., B. Sivyer, and W.R. Taylor, Direction selectivity in the retina: symmetry and asymmetry in structure and function. Nat Rev Neurosci, 2012. 13(3): p. 194–208.

12. Yonehara, K., et al., Identification of retinal ganglion cells and their projections involved in central transmission of information about upward and downward image motion. PLoS One, 2009. 4(1): p. e4320.

13. Yonehara, K., et al., Expression of SPIG1 reveals development of a retinal ganglion cell subtype projecting to the medial terminal nucleus in the mouse. PLoS One, 2008. 3(2): p. e1533.

14. Sun, L.O., et al., Functional assembly of accessory optic system circuitry critical for compensatory eye movements. Neuron, 2015. 86(4): p. 971–984.

15. Osterhout, J.A., et al., Contactin-4 mediates axon-target specificity and functional development of the accessory optic system. Neuron, 2015. 86(4): p. 985–999.

16. Luo, L. and D.D. O’Leary, Axon retraction and degeneration in development and disease. Annu Rev Neurosci, 2005. 28: p. 127–56.

17. Huberman, A.D., M.B. Feller, and B. Chapman, Mechanisms underlying development of visual maps and receptive fields. Annu Rev Neurosci, 2008. 31: p. 479–509.

18. Riccomagno, M.M. and A.L. Kolodkin, Sculpting neural circuits by axon and dendrite pruning. Annu Rev Cell Dev Biol, 2015. 31: p. 779–805.

19. Al-Khindi, T., et al., The transcription factor Tbx5 regulates direction-selective retinal ganglion cell development and image stabilization. Curr Biol, 2022. 32(19): p. 4286–4298.e5.

20. Lin, T.-H., et al., Protein tyrosine phosphatase receptor type kappa (PTPRκ) regulates Superior ON-Direction Selective Ganglion Cell development, facilitating image stabilization. bioRxiv, 2026: p. 2026.01.07.698158.

21. Rheaume, B.A., et al., Single cell transcriptome profiling of retinal ganglion cells identifies cellular subtypes. Nat Commun, 2018. 9(1): p. 2759.

22. Woo, S.H., et al., Piezo2 is required for Merkel-cell mechanotransduction. Nature, 2014. 509(7502): p. 622–6.

23. Toma, K., et al., Perivascular neurons instruct 3D vascular lattice formation via neurovascular contact. Cell, 2024. 187(11): p. 2767–2784.e23.

24. Rowan, S. and C.L. Cepko, Genetic analysis of the homeodomain transcription factor Chx10 in the retina using a novel multifunctional BAC transgenic mouse reporter. Dev Biol, 2004. 271(2): p. 388–402.

25. Nakhai, H., et al., Ptf1a is essential for the differentiation of GABAergic and glycinergic amacrine cells and horizontal cells in the mouse retina. Development, 2007. 134(6): p. 1151–60.

26. Ma, S., et al., Excessive mechanotransduction in sensory neurons causes joint contractures. Science, 2023. 379(6628): p. 201–206.

27. Guo, L., et al., Inducible Rbpms-CreER(T2) Mouse Line for Studying Gene Function in Retinal Ganglion Cell Physiology and Disease. Cells, 2023. 12(15).

28. Kirkby, L.A., et al., A role for correlated spontaneous activity in the assembly of neural circuits. Neuron, 2013. 80(5): p. 1129–44.

29. Malin, J. and C. Desplan, Neural specification, targeting, and circuit formation during visual system assembly. Proc Natl Acad Sci U S A, 2021. 118(28).

30. Lilley, B.N., et al., Genetic access to neurons in the accessory optic system reveals a role for Sema6A in midbrain circuitry mediating motion perception. J Comp Neurol, 2019. 527(1): p. 282–296.

31. Osterhout, J.A., et al., Birthdate and outgrowth timing predict cellular mechanisms of axon target matching in the developing visual pathway. Cell Rep, 2014. 8(4): p. 1006–17.

32. Hong, Y.K., et al., Refinement of the retinogeniculate synapse by bouton clustering. Neuron, 2014. 84(2): p. 332–9.

33. Arroyo, D.A. and M.B. Feller, Spatiotemporal Features of Retinal Waves Instruct the Wiring of the Visual Circuitry. Front Neural Circuits, 2016. 10: p. 54.

34. Kerschensteiner, D., Spontaneous Network Activity and Synaptic Development. Neuroscientist, 2014. 20(3): p. 272–90.

35. Ranade, S.S., et al., Piezo2 is the major transducer of mechanical forces for touch sensation in mice. Nature, 2014. 516(7529): p. 121–5.

36. Woo, S.H., et al., Piezo2 is the principal mechanotransduction channel for proprioception. Nat Neurosci, 2015. 18(12): p. 1756–62.

37. Nonomura, K., et al., Piezo2 senses airway stretch and mediates lung inflation-induced apnoea. Nature, 2017. 541(7636): p. 176–181.

38. Zeng, W.Z., et al., PIEZOs mediate neuronal sensing of blood pressure and the baroreceptor reflex. Science, 2018. 362(6413): p. 464–467.

39. Pampols-Perez, M., et al., Mechanosensitive PIEZO2 channels shape coronary artery development. Nat Cardiovasc Res, 2025. 4(7): p. 921–937.

40. Marshall, K.L., et al., PIEZO2 in sensory neurons and urothelial cells coordinates urination. Nature, 2020. 588(7837): p. 290–295.

41. Hill, R.Z., et al., Renal PIEZO2 is an essential regulator of renin. Cell, 2026. 189(1): p. 161–178.e22.

42. Wang, Y., et al., A key role of PIEZO2 mechanosensitive ion channel in adipose sensory innervation. Cell Metab, 2025. 37(4): p. 1001–1011.e7.

43. Koser, D.E., et al., Mechanosensing is critical for axon growth in the developing brain. Nat Neurosci, 2016. 19(12): p. 1592–1598.

44. Song, Y., et al., The Mechanosensitive Ion Channel Piezo Inhibits Axon Regeneration. Neuron, 2019. 102(2): p. 373–389.e6.

45. Tao, L., et al., Dendrites use mechanosensitive channels to proofread ligand-mediated neurite extension during morphogenesis. Dev Cell, 2022. 57(13): p. 1615–1629.e3.

46. Xie, Q., et al., Transcription factor Acj6 controls dendrite targeting via a combinatorial cell-surface code. Neuron, 2022. 110(14): p. 2299–2314.e8.

47. Delle Vedove, A., et al., Biallelic Loss of Proprioception-Related PIEZO2 Causes Muscular Atrophy with Perinatal Respiratory Distress, Arthrogryposis, and Scoliosis. Am J Hum Genet, 2016. 99(5): p. 1206–1216.

48. Mulhall, E.M., et al., The molecular basis of force selectivity by PIEZO2. Nature, 2026. 653(8113): p. 297–305.

49. Pardo-Pastor, C., et al., Piezo2 channel regulates RhoA and actin cytoskeleton to promote cell mechanobiological responses. Proc Natl Acad Sci U S A, 2018. 115(8): p. 1925–1930.

50. Ghose, A. and P. Pullarkat, The role of mechanics in axonal stability and development. Semin Cell Dev Biol, 2023. 140: p. 22–34.

51. Anava, S., et al., The regulative role of neurite mechanical tension in network development. Biophys J, 2009. 96(4): p. 1661–70.

52. Gomez, T.M. and N.C. Spitzer, In vivo regulation of axon extension and pathfinding by growth-cone calcium transients. Nature, 1999. 397(6717): p. 350–5.

53. Carrillo, R.A., et al., Presynaptic activity and CaMKII modulate retrograde semaphorin signaling and synaptic refinement. Neuron, 2010. 68(1): p. 32–44.

54. Santiago, C., et al., Activity-dependent development of the body’s touch receptors. Neuron, 2025. 113(11): p. 1758–1773.e9.

55. Zhang, Y., et al., PIEZO channels link mechanical forces to uterine contractions in parturition. Science, 2025. 390(6774): p. eady3045.

56. Coste, B., et al., Gain-of-function mutations in the mechanically activated ion channel PIEZO2 cause a subtype of Distal Arthrogryposis. Proc Natl Acad Sci U S A, 2013. 110(12): p. 4667–72.

57. McMillin, M.J., et al., Mutations in PIEZO2 cause Gordon syndrome, Marden-Walker syndrome, and distal arthrogryposis type 5. Am J Hum Genet, 2014. 94(5): p. 734–44.

58. Akinci, G., et al., Genetic and clinical spectrum of PIEZO2-related disorders: insights from a multicenter study of 26 patients. Neuromuscul Disord, 2025. 53: p. 105423.

59. Beals, R.K. and R.G. Weleber, Distal arthrogryposis 5: a dominant syndrome of peripheral contractures and ophthalmoplegia. Am J Med Genet A, 2004. 131(1): p. 67–70.

60. Oystreck, D.T., Ophthalmoplegia and Congenital Cranial Dysinnervation Disorders. J Binocul Vis Ocul Motil, 2018. 68(1): p. 31–33.

61. Kiraly, J.K., et al., Slit2/Robo1 signaling constrains image stabilization responses to preserve ethologically favorable directional asymmetry. Curr Biol, 2025. 35(19): p. 4714–4726.e4.

62. Leinninger, G.M., et al., Leptin action via neurotensin neurons controls orexin, the mesolimbic dopamine system and energy balance. Cell Metab, 2011. 14(3): p. 313–23.

63. Stahl, J.S., Using eye movements to assess brain function in mice. Vision Res, 2004. 44(28): p. 3401–10.

64. Picelli, S., et al., Full-length RNA-seq from single cells using Smart-seq2. Nat Protoc, 2014. 9(1): p. 171–81.

65. Kim, D., B. Langmead, and S.L. Salzberg, HISAT: a fast spliced aligner with low memory requirements. Nat Methods, 2015. 12(4): p. 357–60.

66. Li, H., et al., The Sequence Alignment/Map format and SAMtools. Bioinformatics, 2009. 25(16): p. 2078–9.

67. Trapnell, C., et al., Differential gene and transcript expression analysis of RNA-seq experiments with TopHat and Cufflinks. Nat Protoc, 2012. 7(3): p. 562–78.

68. Butler, A., et al., Integrating single-cell transcriptomic data across different conditions, technologies, and species. Nat Biotechnol, 2018. 36(5): p. 411–420.

69. Stuart, T., et al., Comprehensive Integration of Single-Cell Data. Cell, 2019. 177(7): p. 1888–1902.e21.

