## Supplemental Information for "The mechanotransduction channel Piezo2 refines axonal projections to the accessory optic system and regulates the optokinetic reflex"

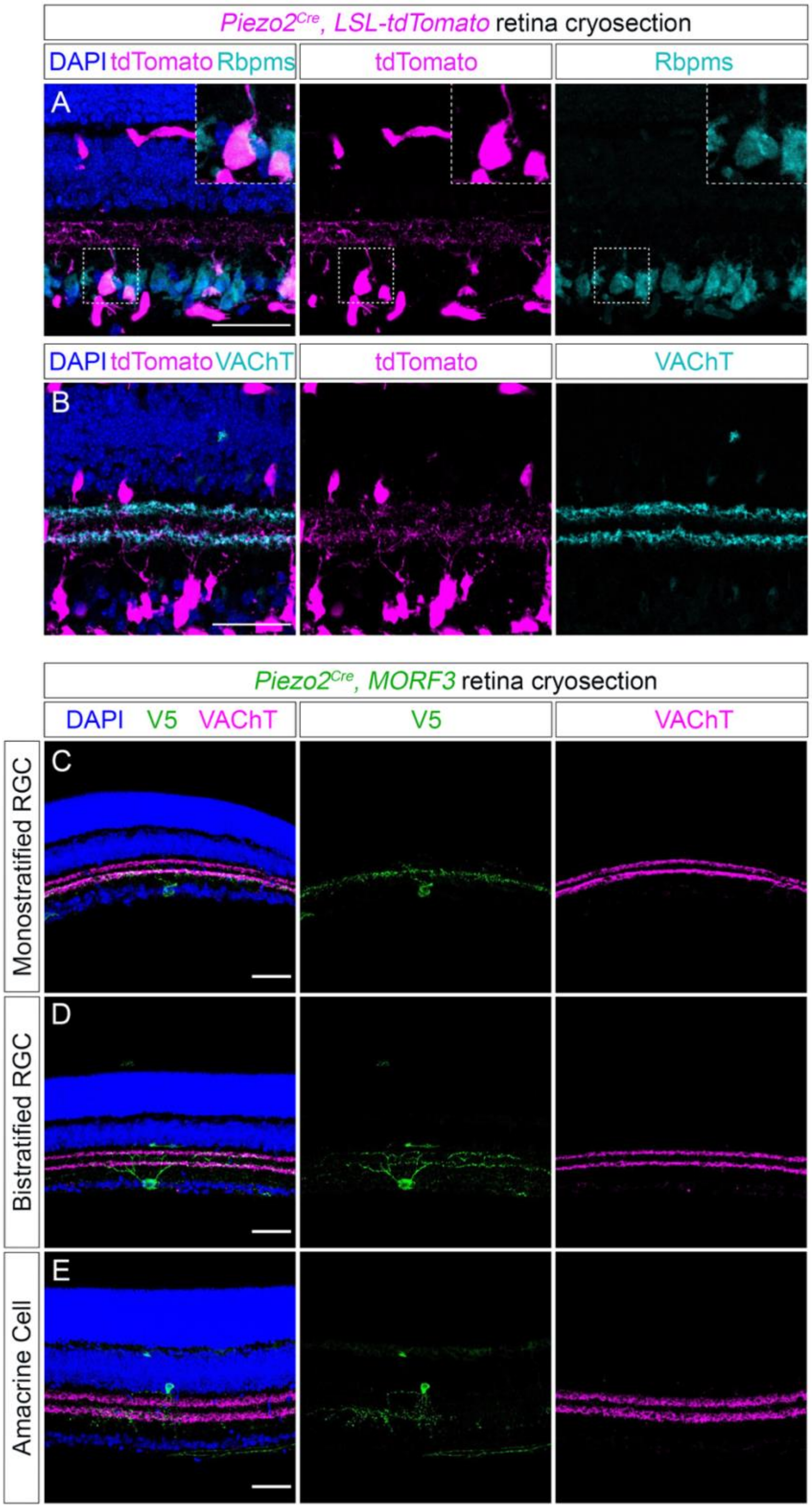


**Figure S1 (Related to Figure 1). Piezo2 is expressed in subsets of non-SAC amacrine cells and RGCs that co-stratify with SACs**

(**A, B**) Retina cryosections from *Piezo2$^{Cre}$; LSL-tdTomato* mice show that *Piezo2* is expressed in a subset of Rbpms$^{+}$ RGCs (**A**). Piezo2$^{+}$ RGCs stratify in S2 and S4 of the IPL, as labeled by VAChT (**B**); *Piezo2* is also expressed in subsets of amacrine cells whose cell bodies reside in the INL, and which do not co-label with VAChT, identifying them as non-SAC amacrine cells (**B**). (**C-E**) Sparse genetic labeling in *Piezo2$^{Cre}$; MORF3* mouse retinas reveals subtypes of ON-monostratified RGCs (**C**), ON-OFF bistratified RGCs (**D**), and amacrine cells (**E**), each expressing a membrane-targeted V5 tag.

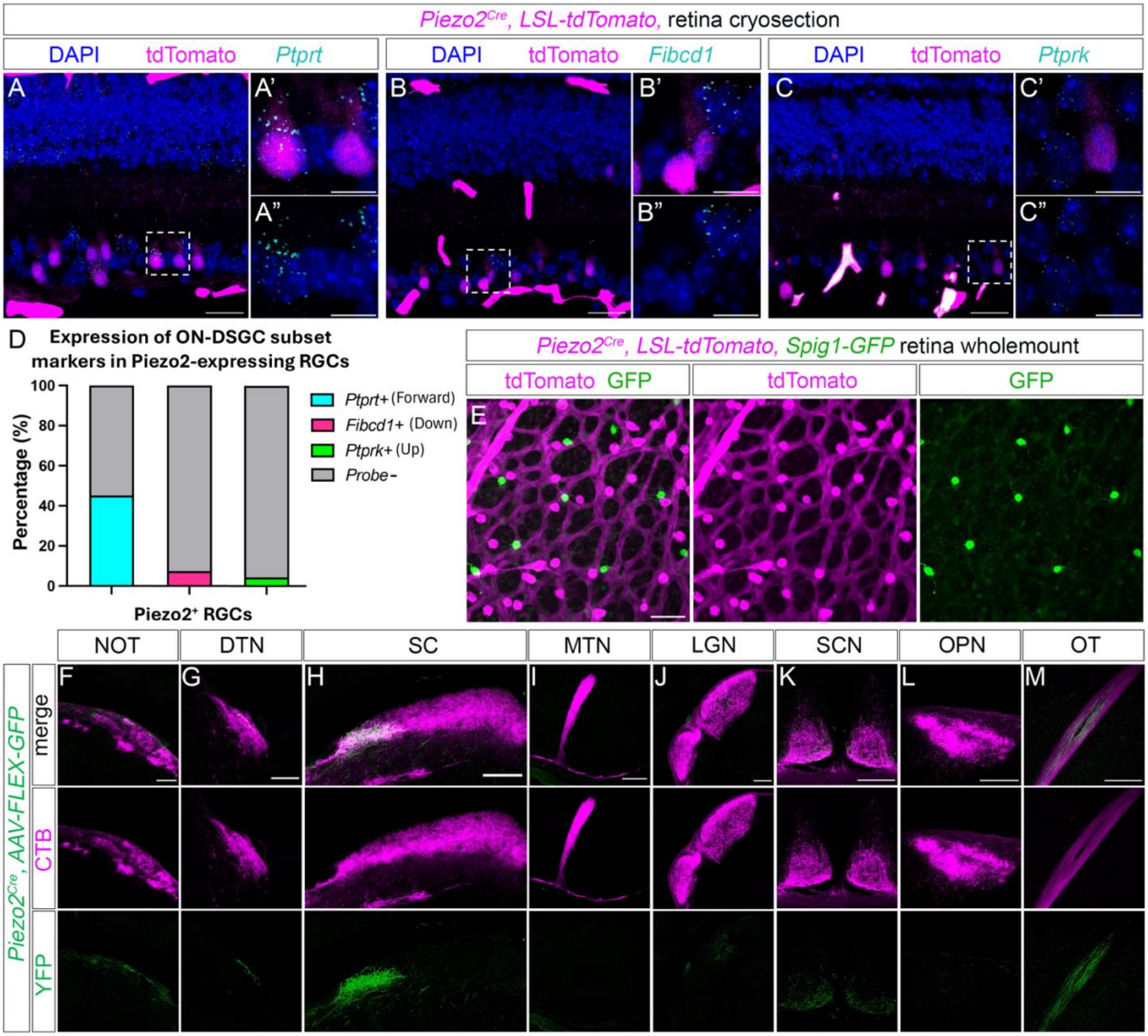


**Figure S2 (related to Figure 2). Piezo2 is not expressed in Up and Down ON-DSGCs**
(**A-C**) RNAscope *in situ* hybridization in *Piezo2$^{Cre}$; LSL-tdTomato* retinal cross sections. *Ptprt* is expressed in a subset of tdTomato$^+$/Piezo2$^+$ cells (**A-A"**); however, neither *Fibcd1* (**B-B"**), nor *Ptprk* (**C-C"**) expression was detected in tdTomato$^+$/Piezo2$^+$ cells. (**D**) Quantification of the percentage of tdTomato$^+$/Piezo2$^+$ cells expressing *Ptprt, Fibcd1*, or *Ptprk*. (**E**) Genetic labeling of Piezo2$^+$ cells in *Piezo2$^{Cre}$; LSL-tdTomato; Spig1-GFP* retinas shows no overlap between tdTomato and GFP signal, confirming that Spig1-GFP$^+$ U-oDSGCs do not express *Piezo2*. (**F-M**) Co-injection of a Cre-dependent AAV encoding GFP (AAV-FLEX-GFP) and CTB-555 at P2 labels Piezo2$^+$ RGC projections and retinorecipient nuclei at P18. Piezo2$^+$ RGCs innervate the NOT (**F**), DTN (**G**), and SC (**H**), but not the MTN (**I**) or the OPN (**L**). Minimal innervation is seen in the LGN (**J**) and SCN (**K**). Piezo2$^+$ RGC axons can be seen traversing the optic tract (**M**).

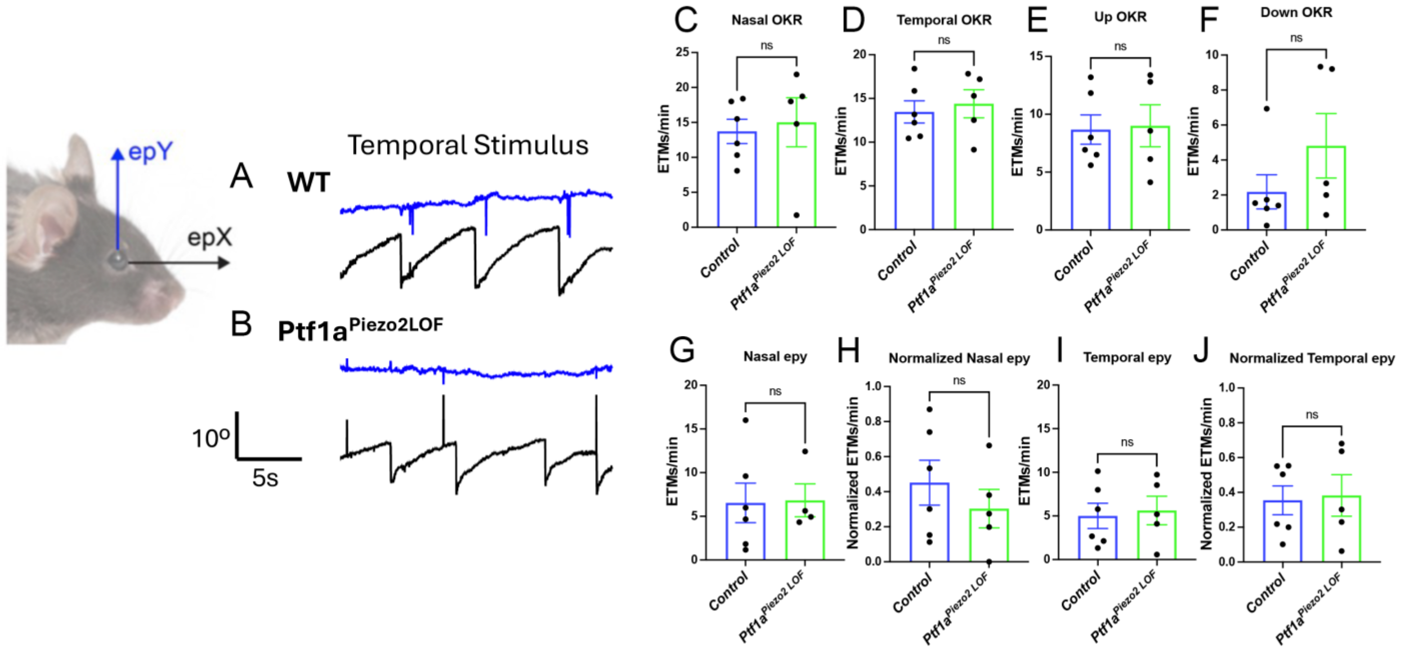


**Figure S3 (related to Figure 4). Piezo2 function in amacrine cells is dispensable for the OKR**
(**A, B**) Amacrine cell-specific Piezo2$^{LOF}$ mice (*Ptf1a*$^{Piezo2LOF}$) do not recapitulate the cross-coupling seen in *Ret*$^{Piezo2LOF}$ mice, showing minimal vertical eye movements in response to temporal stimuli (**B**). (**C-F**) *Ptf1a*$^{Piezo2LOF}$ mice show normal OKR responses to continuous gratings moving in the nasal (**C**), temporal (**D**), upward (**E**), or downward (**F**) direction. (**G-J**) *Ptf1a*$^{Piezo2LOF}$ mice show no statistically significant differences in absolute and normalized number of in-phase upward vertical ETMs/min (epY) during nasal (**G, H**) or temporal (**I, J**) stimuli compared to WT mice.
